# MicroRNA correlates of resilience to Alzheimer’s disease identify candidate therapeutic targets for neuroprotection

**DOI:** 10.64898/2026.08.04.737285

**Authors:** Pourya Naderi, Latika Matai, Quadri Adewale, Isabel Castanho, Alice Rodrigues, Maria Mavrikaki, Ioannis S. Vlachos, Frank J. Slack, Winston Hide

**Author notes:** Corresponding author(s) contact details: Frank Slack; Winston Hide. Equally contributing senior authors (Frank J. Slack, Winston Hide).

## Abstract

Many aged individuals accumulate advanced Alzheimer’s disease (AD) neuropathology without cognitive decline, signifying potent endogenous resilience mechanisms. Mimicking resilience to AD may offer new therapeutic opportunities by emulating endogenous neuroprotection, but the molecular pathways that underpin resilience remain largely unknown. MicroRNAs (miRNAs)—small noncoding RNA molecules that post-transcriptionally regulate gene expression—influence neuronal and glial processes associated with development, aging, and AD neurodegeneration but their role in resilience has not been characterized. We investigated resilience-associated miRNAs and their targeted programs in the dorsolateral prefrontal cortex of post-mortem human brains. We analyzed matched miRNA and messenger RNA (mRNA) expression across resilient, AD, and control subjects from the Religious Order Study and Memory and Aging Project (ROSMAP) cohort. Nine miRNAs were differentially expressed in resilience compared with AD, including previously unreported miR-362-3p and miR-433-3p and new resilience-associated roles for known AD-related miRNAs, including neuronal miR-132-3p and miR-129-5p. By analyzing miRNA activity across AD progression, we found additional cognitive-associated miRNAs (e.g., miR-335-5p and miR-19b) and one plaque-restricted miRNA (miR-199a-5p). Sex-specific miRNA dysregulation was observed: miR-7-5p showed male-specific upregulation in AD versus resilience and suggested sex-specific differences in AD patients. Integrated co-expression and target-enrichment analyses linked resilience-associated miRNAs to pathways that were associated with cognitive decline, including transforming growth factor β, Rho guanosine triphosphatases, and neurotransmitter receptor regulation. We also report restricted co-activity in AD subjects for miR-362-3p with inflammatory pathways, including tumor necrosis factor and Toll-like receptor signaling. These results demonstrate systematic involvement of miRNAs across neuronal and glial programs of AD resilience, cognitive decline, and sex-specific regulation. Our study provides an important resource for discovery of actionable regulatory programs that could lead to new therapies, based on endogenous molecules, that emulate natural resilience to AD.

**Key points:**

- Nine distinct cortical microRNA (miRNA) signatures correlate with resilience to Alzheimer’s disease (AD)-related cognitive decline, pointing to endogenous regulatory programs that may help preserve cognitive function.
- At the miRNA level, resilient brains show minimal or no detectable differences to healthy individuals, despite the profound pathological differences, supporting the idea that resilience reflects preserved molecular homeostasis rather than a separate disease state.
- Some miRNA signatures are specific to a certain pathology (e.g., one linked to amyloid-β plaques independently of tau pathology and cognitive decline), helping to disentangle the regulation of these processes.
- Sex-specific miRNA dysregulation in AD suggests that resilience-linked pathways may be regulated differently in males and females, reinforcing the value of sex-stratified analyses and the potential for differential therapeutic strategies.
- Integrative miRNA–pathway analysis highlights candidate regulatory networks involved in inflammation, matrisome, and cellular stress responses, providing a framework for resilience-linked therapeutic targets.

## Introduction

Alzheimer’s disease (AD) is biologically and clinically defined based on unique neuropathological findings (Jack et al., 2024) and progressive cognitive loss that leads to dementia. Symptomatic treatments aimed at modifying pathology or inflammation, such as cholinesterase inhibitors and emerging immunotherapies targeting amyloid-β (Aβ), provide limited benefit and have not yielded substantial clinical success (Long and Holtzman, 2019; Aljuhani et al., 2024; Jin et al., 2024; Zuliani et al., 2024). New therapies that directly target neuroprotection as an alternative to symptomatic treatment of late-stage pathology are urgently needed. A considerable portion of the aging population lives without cognitive loss while showing advanced AD pathology at autopsy, a phenomenon referred to as resilience to AD (Gómez-Isla and Frosch, 2022; Montine et al., 2023; Castanho et al., 2025). Longitudinal community and clinical cohorts have reported that between 10 to 30% of participants with healthy cognition met the pathological criteria for AD (Riley et al., 2005; Schneider et al., 2009; Corrada et al., 2012; Gómez-Isla and Frosch, 2022). A recent large-scale population study of AD biomarkers in Norway has revealed that approximately 20% of individuals aged 70+ have AD neuropathology with either no or mild cognitive impairment (Aarsland et al., 2025). The existence of potent endogenous resilience mechanisms that prevent the onset of AD dementia provides a unique opportunity to discover novel therapies. Interventions that emulate endogenous resilience do not exist yet, partly due to limited understanding of actionable molecular programs underlying resilience and protection.

Resilience against AD is multifaceted and includes balancing excitatory/inhibitory neuronal populations (Castanho et al., 2025), effective clearance of Aβ plaques, and suppression of neuroinflammation (Gómez-Isla and Frosch, 2022). Recent studies show that resilience incorporates molecular and cellular events at a transcriptomic level (Perez-Nievas et al., 2013; Pedrinolla et al., 2017; Ridge et al., 2017; Mostafavi et al., 2018; Dumitrescu et al., 2020; O’Neill et al., 2024; Castanho et al., 2025). Most of these studies, however, primarily focus on protein-coding genes, which comprise only a fraction of the transcriptome. In contrast, few studies have explored non-coding molecular factors of resilience, even though non-coding RNAs represent the majority of the human transcriptome (Mattick, 2001; Kelley et al., 2024; Counts et al., 2025). As such, the roles of non-coding RNAs in regulation of resilience to AD remain unknown.

MicroRNAs (miRNAs) are a conserved class of non-coding RNAs that play a central role in post-transcriptional regulation. These short RNAs (∼21 nucleotides) repress gene expression by binding to the 3’ untranslated regions (3’ UTRs) of target messenger RNAs (mRNAs), leading to RNA degradation and/or translational suppression. miRNAs regulate key associated mechanisms of AD including aging, senescence, and Aβ and tau pathologies (Bartel, 2004; Thalyana and Slack, 2012; Lau et al., 2013; Suh, 2018; Samadian et al., 2021; Li et al., 2024; Polzer et al., 2025). miRNAs influence gene networks involved in neurogenesis, synaptic integrity, microglial activation, and brain metabolic homeostasis, and their dysregulation has been consistently linked to Aβ, tau, and neuroinflammatory pathways in AD (Hébert et al., 2008; Absalon et al., 2013; Banzhaf-Strathmann et al., 2014; Kaur et al., 2023; Yin et al., 2023). For example, miR-132-3p regulates tau phosphorylation through the acetyltransferase EP300 and glycogen synthase kinase-3 β (GSK3B), modulates adult hippocampal neurogenesis, and is downregulated in the brain during AD progression (Wang et al., 2017; El Fatimy et al., 2018). miR-155 modulates Aβ clearance by regulating disease-associated microglia in AD and is being evaluated in clinical trials for therapeutics (Yin et al., 2023). miRNAs are strong modulators of central nervous system (CNS) cell states and can potentially program cells towards phenotypes associated with resilience. For example, miR-9-3p/5p and miR-124-3p induce neuronal fate and fibroblast-to-neuron transformation *in vitro* (Cates et al., 2021; Church et al., 2021; Sun et al., 2024). Collectively, evidence points to the involvement of miRNAs across a broad range of brain cell types and biological processes involved in AD, including microglial activation, neurogenesis, and tau pathology, making them valuable targets for intervention. However, the roles of miRNAs in resilience and their broader mechanisms in AD pathogenesis remain poorly understood.

We systematically explored miRNAs associated with human AD resilience by analyzing publicly available bulk gene and miRNA expression data from the dorsolateral prefrontal cortex (DLPFC) of post-mortem AD and unaffected brains from the Religious Orders Study Memory and Aging Project (ROSMAP) cohort (Patrick et al., 2017; De Jager et al., 2018). The DLPFC represents a highly affected brain region in AD patients, and it has been profiled across hundreds of participants in the ROSMAP multiomic cohort (De Jager et al., 2018). We analyzed concordant miRNA/mRNA expression profiles, focusing on resilience, and identified miRNAs uniquely associated with resilience, Aβ burden, cognitive performance, and sex-specific pathogenesis. By leveraging systems biology methods, we moved beyond analysis of isolated miRNA–mRNA interaction events to examine global dysregulation of transcriptional networks and the activation of intracellular signaling pathways governed by distinct regulatory miRNAs.

## Results

### Dataset preparation and sample classification

To investigate the interplay between miRNAs and gene expression, we analyzed publicly available miRNA (*n* = 568) and gene (*n* = 631) expression data from post-mortem DLPFC tissues from aged individuals spanning the AD spectrum (**Figure 1A**), obtained from the ROSMAP cohort (De Jager et al., 2018; Patrick et al., 2017). Subjects with miRNA expression (derived using NanoString nCounter) were classified into AD (*n* = 162), resilient (*n* = 63), and control (*n* = 43) groups based on levels of pathology and cognitive decline (**Table 1**), following criteria from our previous study (Castanho et al., 2025). Briefly: AD subjects had high levels of Aβ plaques (frequent or moderate neuritic plaques) and tau pathology (Braak Stage ³ III) along with a diagnosis of AD dementia (with no other forms of dementia); resilient subjects had high levels of plaques and tangles with no cognitive impairment; and control subjects had low levels of plaques and tangles along with no cognitive impairment (see Methods). An additional category of participants with presymptomatic AD, characterized by the high levels of pathologies seen in resilience and AD but with mild cognitive impairment (MCI), was defined to further characterize resilience with respect to MCI. For downstream co-expression analysis, we also leveraged a subset of 475 ROSMAP profiles with both miRNA and mRNA expression data that had a cognitive assessment of either healthy (*n* = 167), MCI (*n* = 126), or AD dementia (*n* = 182), regardless of high or low levels of Aβ and tau pathologies (Patrick et al., 2017; De Jager et al., 2018). This sample set offered the advantage of incorporating additional participants that did not fit into any of the categories listed above (e.g., high plaques and low tangles, or MCI with no pathology).

**Figure 1.**
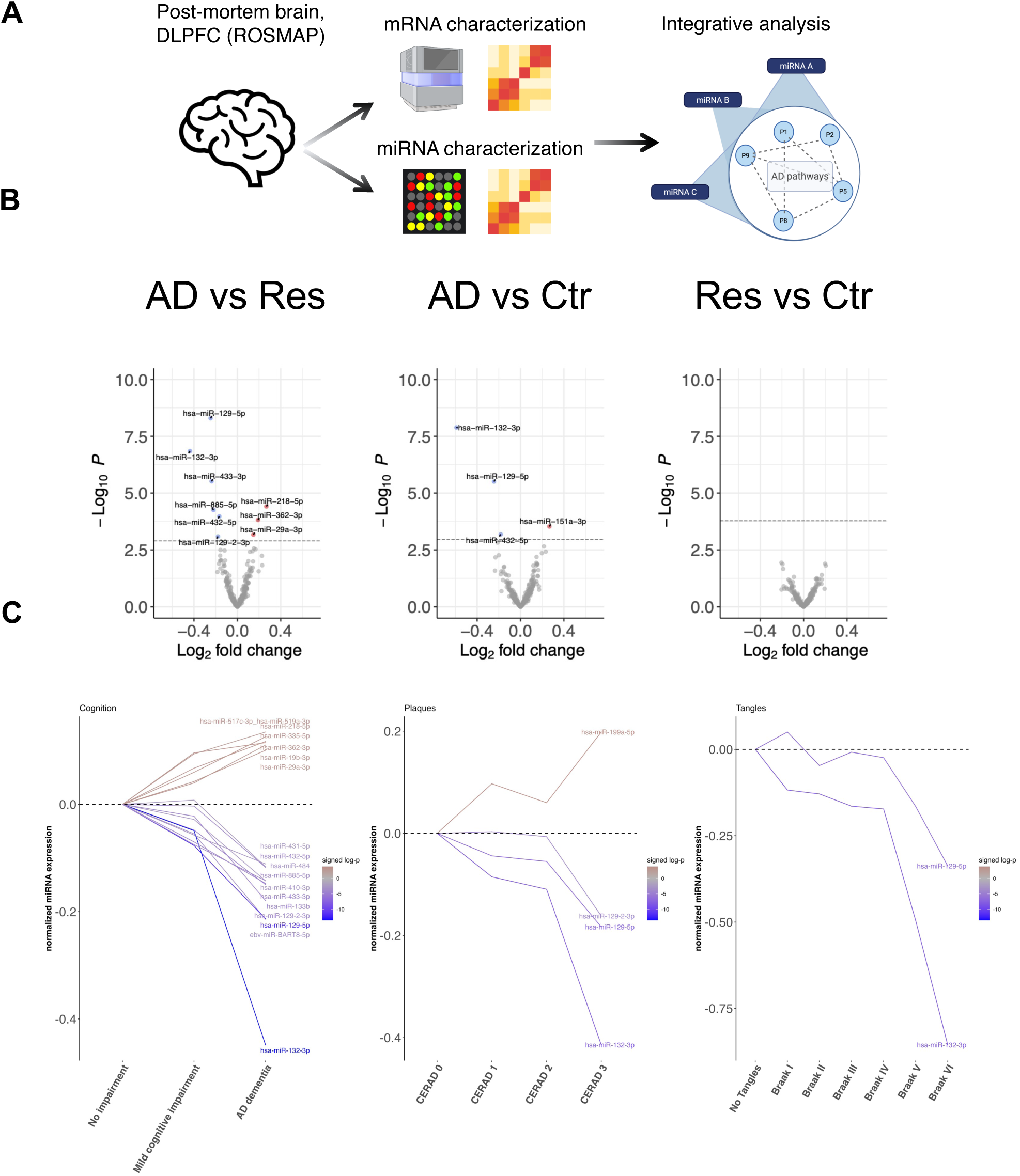
miRNA expression analysis. **A.** study overview. **B.** Differential expression analysis between AD, Resilience (Res), and Control (Ctr) groups. Differential expression was performed using a linear model accounting for covariates and assessed using a two-sided moderated t-test, implemented in the Limma package. Colored points represent significant differential expression (Benjamini-Hochberg false discovery rate (FDR) < 0.05). Dashed lines in each panel represent the p-value corresponding to FDR < 0.05 cut-off. **C.** Panels show miRNAs with statistically significant trends associated with the progression of cognitive decline and/or AD pathology. Y-axes represent the mean of normalized miRNA expression across all samples corresponding to each value on the Y-axis, centered to the lowest category on the Y-axis (e.g. no tangles). Significant trends were determined using a proportional odds model for ordinal categorical regression. The model was implemented with a cumulative logit link using the VGAM package. P-values were derived from two-sided z-scores of the proportional odds model.

**Table 1.**
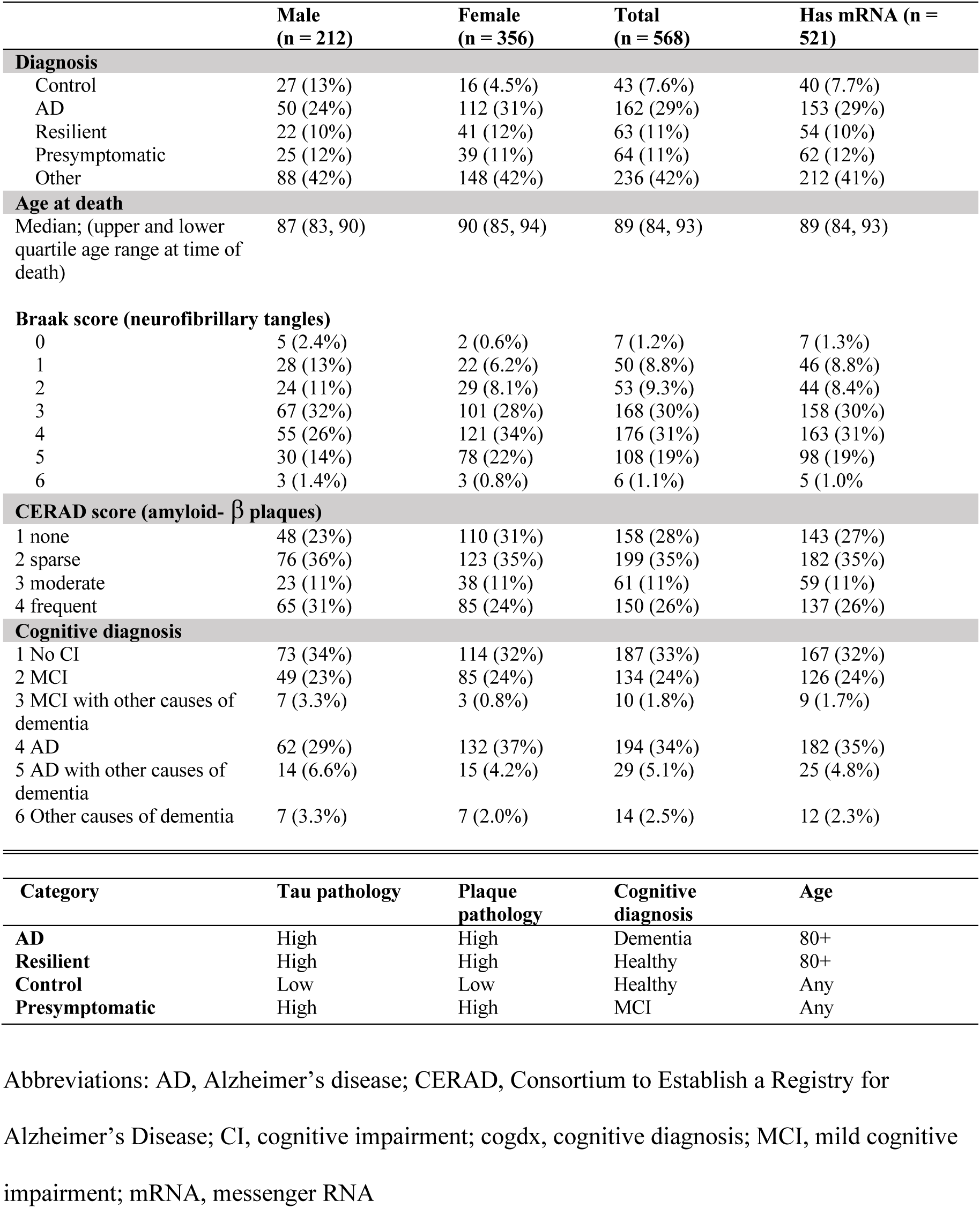
Summary statistics of microRNA (miRNA) expression dataset.

|  | Male<br>(n = 212) | Female<br>(n = 356) | Total<br>(n = 568) | Has mRNA (n = 521) |
| --- | --- | --- | --- | --- |
| **Diagnosis** |  |  |  |  |
| Control | 27 (13%) | 16 (4.5%) | 43 (7.6%) | 40 (7.7%) |
| AD | 50 (24%) | 112 (31%) | 162 (29%) | 153 (29%) |
| Resilient | 22 (10%) | 41 (12%) | 63 (11%) | 54 (10%) |
| Presymptomatic | 25 (12%) | 39 (11%) | 64 (11%) | 62 (12%) |
| Other | 88 (42%) | 148 (42%) | 236 (42%) | 212 (41%) |
| **Age at death** |  |  |  |  |
| Median; (upper and lower quartile age range at time of death) | 87 (83, 90) | 90 (85, 94) | 89 (84, 93) | 89 (84, 93) |
| **Braak score (neurofibrillary tangles)** |  |  |  |  |
| 0 | 5 (2.4%) | 2 (0.6%) | 7 (1.2%) | 7 (1.3%) |
| 1 | 28 (13%) | 22 (6.2%) | 50 (8.8%) | 46 (8.8%) |
| 2 | 24 (11%) | 29 (8.1%) | 53 (9.3%) | 44 (8.4%) |
| 3 | 67 (32%) | 101 (28%) | 168 (30%) | 158 (30%) |
| 4 | 55 (26%) | 121 (34%) | 176 (31%) | 163 (31%) |
| 5 | 30 (14%) | 78 (22%) | 108 (19%) | 98 (19%) |
| 6 | 3 (1.4%) | 3 (0.8%) | 6 (1.1%) | 5 (1.0%) |
| **CERAD score (amyloid- $\beta$ plaques)** |  |  |  |  |
| 1 none | 48 (23%) | 110 (31%) | 158 (28%) | 143 (27%) |
| 2 sparse | 76 (36%) | 123 (35%) | 199 (35%) | 182 (35%) |
| 3 moderate | 23 (11%) | 38 (11%) | 61 (11%) | 59 (11%) |
| 4 frequent | 65 (31%) | 85 (24%) | 150 (26%) | 137 (26%) |
| **Cognitive diagnosis** |  |  |  |  |
| 1 No CI | 73 (34%) | 114 (32%) | 187 (33%) | 167 (32%) |
| 2 MCI | 49 (23%) | 85 (24%) | 134 (24%) | 126 (24%) |
| 3 MCI with other causes of dementia | 7 (3.3%) | 3 (0.8%) | 10 (1.8%) | 9 (1.7%) |
| 4 AD | 62 (29%) | 132 (37%) | 194 (34%) | 182 (35%) |
| 5 AD with other causes of dementia | 14 (6.6%) | 15 (4.2%) | 29 (5.1%) | 25 (4.8%) |
| 6 Other causes of dementia | 7 (3.3%) | 7 (2.0%) | 14 (2.5%) | 12 (2.3%) |
| **Category** | **Tau pathology** | **Plaque pathology** | **Cognitive diagnosis** | **Age** |
| AD | High | High | Dementia | 80+ |
| Resilient | High | High | Healthy | 80+ |
| Control | Low | Low | Healthy | Any |
| Presymptomatic | High | High | MCI | Any |

Abbreviations: AD, Alzheimer's disease; CERAD, Consortium to Establish a Registry for Alzheimer's Disease; CI, cognitive impairment; cogdx, cognitive diagnosis; MCI, mild cognitive impairment; mRNA, messenger RNA

### Resilience is marked by distinct associated miRNAs

To discover miRNAs associated with resilience and compare them with other AD-associated miRNAs, we compared miRNA expression between AD, resilient, and control subjects. In the comparison between AD and resilience, nine miRNAs showed statistically significant differential expression (adjusted *p* <0.05) after accounting for sex, age, post-mortem interval, apolipoprotein E *(APOE)* genotype, and RNA integrity number (**Figure 1B**). Differentially expressed miRNAs in AD versus resilience included some with little or no known direct link to AD (upregulated: miR-362-3p and miR-218-5p; downregulated: miR-433-3p, miR-432-5p, and miR-885-5p). We also observed miRNAs with differential expression in AD versus resilience that were previously associated with AD or neuroprotection (upregulated: miR-29a-3p; downregulated: miR-132-3p and miR-129-5p, and miR-129-2-3p). In the AD versus control comparison, we observed significant downregulation of miR-132-3p and miR-129-5p along with novel findings of downregulation of miR-432-5p and miR-151a-3p. Notably, comparison of resilient and control subjects yielded no significant differentially expressed miRNAs, consistent with our previous transcriptomic analysis of protein-coding genes, which also showed minimal differences between resilient and control subjects (Castanho et al., 2025). The following comparisons also did not yield any significant miRNAs: presymptomatic versus control and resilient versus presymptomatic (**Supplementary Table S1**).

In summary, multiple miRNAs were downregulated in AD compared with both resilient and control subjects (miR-132-3p, miR-129-5p, and miR-432-5p). However, other miRNAs showed AD-associated differences relative to only one comparison group—either controls (miR-151a-3p) or resilient subjects (miR-362-3p)—indicating both shared and group-specific expression patterns.

### miRNA expression tracks progression of AD pathology and cognition loss

AD is defined across a spectrum of clinical (cognitive) and pathological burden. To determine miRNAs associated with AD progression (i.e., cognitive-associated, plaque-associated, and tangle-associated), we employed an ordinal categorical linear model to examine the relationships between miRNA expression and cognitive status, Aβ pathology, and tau pathology. This approach, known as the proportional odds model, is a multi-level extension of logistic regression and offers the advantage of using a larger sample size than standard differential expression analysis between binarized groups.

miR-132-3p and miR-129-5p significantly tracked the burden of cognitive decline, Aβ plaques, and tau pathology (**Figure 1C)**. Cognitive-associated miRNAs included all differentially expressed miRNAs in AD versus resilience as well as additional significant candidates not captured as significant via differential expression analysis, including miR-133b and ebv-miR-BART8-5p (**Supplementary Table S2**). While we primarily focus on human miRNAs, it is noteworthy that the Nanostring nCounter assay used to generate the miRNA profiles contains probes for viral-encoded miRNAs, including ebv-miR-BART8-5p, which is an Epstein-Barr virus (EBV) miRNA. While EBV infection is prevalent in human populations (Zhang et al., 2021), the significance of these findings remains to be tested. We also identified a unique significant association between miR-199a-5p and Aβ pathology, with no significant association with tangles or cognitive decline, nor differential expression between any two categories. Our results demonstrate that miRNAs can reflect and delineate disease progression across specific clinical and pathological features of AD.

### miRNAs capture sex-specific events in AD

We investigated differentially expressed miRNAs between male and female subjects across AD, resilient, and control conditions, using a linear model with interaction terms between the covariates encoding sex and diagnosis. We determined both significant [false discovery rate (FDR) <0.05] and suggestive (*p* <0.05 and FDR <0.2) miRNAs across comparisons and compared findings with differential expression analysis and pathology associations across all subjects (**Supplementary Table S3**). miR-7-5p showed significant male-specific upregulation in AD versus resilience without significant differential expression in females or in all subjects (**Figure 2**). This miRNA also showed suggestive differential expression between male and female AD subjects. The dysregulation of miR-7-5p between AD versus resilient subjects was higher in males than in females. Also, miR-218-5p showed significant upregulation in AD versus resilience only in male subjects. Nine miRNAs had suggestive (*p* <0.05, FDR <0.2) dysregulation between AD males and AD females, including miR-150-5p, miR-199b-5p, miR-34b-3p, miR-361-5p, miR-30c-5p, miR-7-5p, miR-328-3p, miR-193b-3p, and miR-26a-5p. None showed concordant significant or suggestive changes associated with AD, resilience, cognition, or pathology across all subjects together. We did not find significant differences in miRNA expression between males and females in either the control or resilience groups.

**Figure 2.**
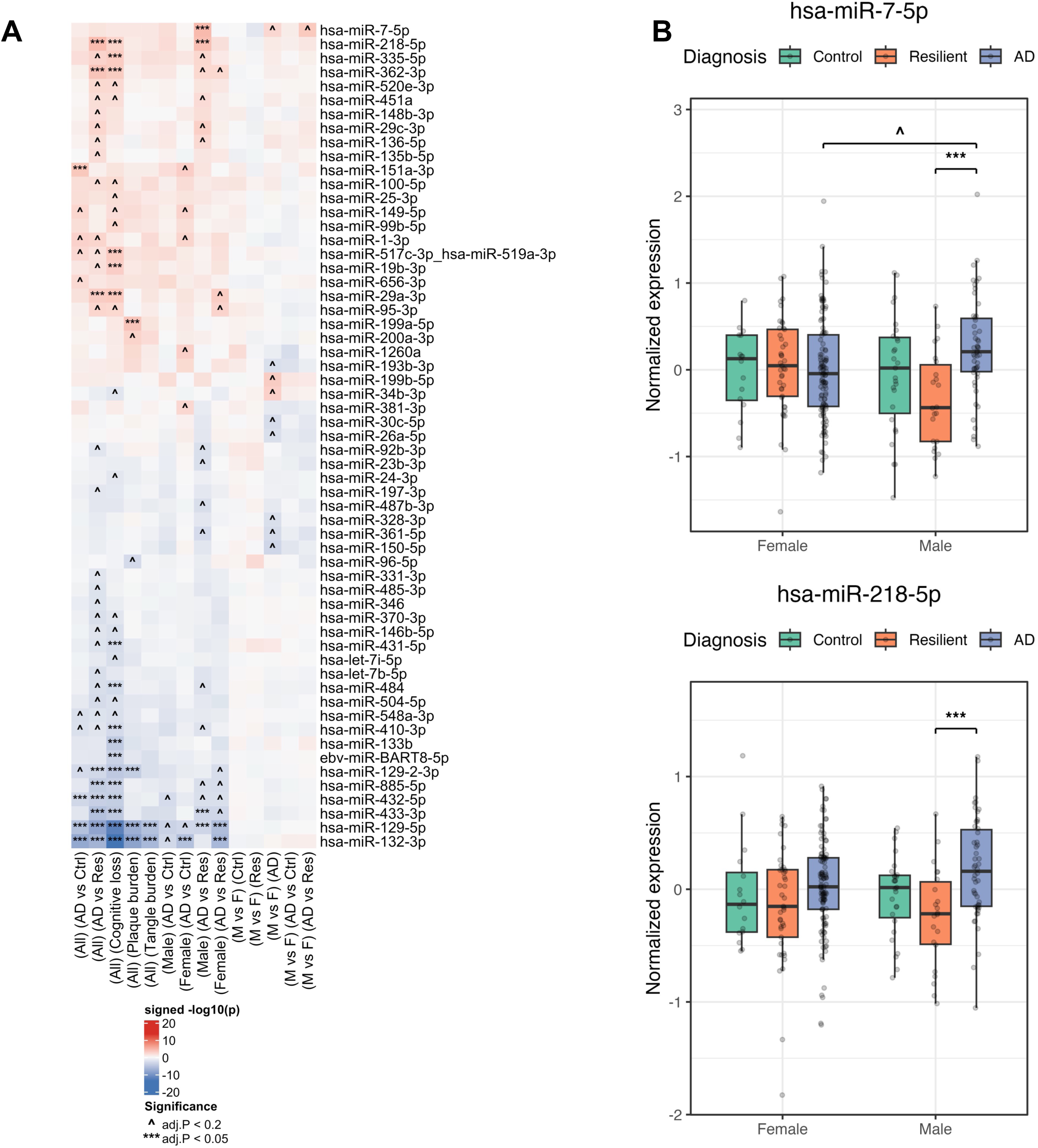
Sex-specific miRNA expression analysis. **A.** Heatmap showing differential expression p-values across varying contrasts. Sex-specific differential expression analysis was performed using a linear model with an interaction term between covariates for sex and diagnosis (AD, Res, Control,…). Each column in the heatmap represents a contrasts, (All: non-sex specific; Male: males only; Female: Females only; M vs F: comparison of males and females). Cell colors represent signed log p-values. P-values were derived using moderated t-test, implemented in the Limma package. P-values were adjusted for multiple comparisons using the Benjamini-Hochberg FDR. ***: FDR < 0.05. ^: FDR < 0.2 and P < 0.05. **B.** Boxplot the levels of miRNAs with sex-specific differential expression. P-values were derived using moderated t-test, implemented in the Limma package. P-values were adjusted for multiple comparisons using the Benjamini-Hochberg FDR. ***: FDR < 0.05. ^: FDR < 0.2 and P < 0.05.

### Integrated miRNA–mRNA co-expression network

To reveal the dynamic interplay between miRNAs and their mRNA targets in AD, we performed co-expression analysis of miRNA and mRNA profiles from the ROSMAP cohort. To recapitulate an unbiased co-activity network and see condition-specific activity, we analyzed co-expression in AD and non-AD subjects, then merged the two into a consensus co-activity. Co-activity of miRNAs and mRNAs were reported when there was evidence of: significant miRNA–mRNA co-expression, miRNA–mRNA binding (retrieved from the TarBase V9 database) (Skoufos et al., 2024), and cognitive association for both miRNA and targeted mRNA (**Figure 3A; Supplementary Table S4**).

**Figure 3.**
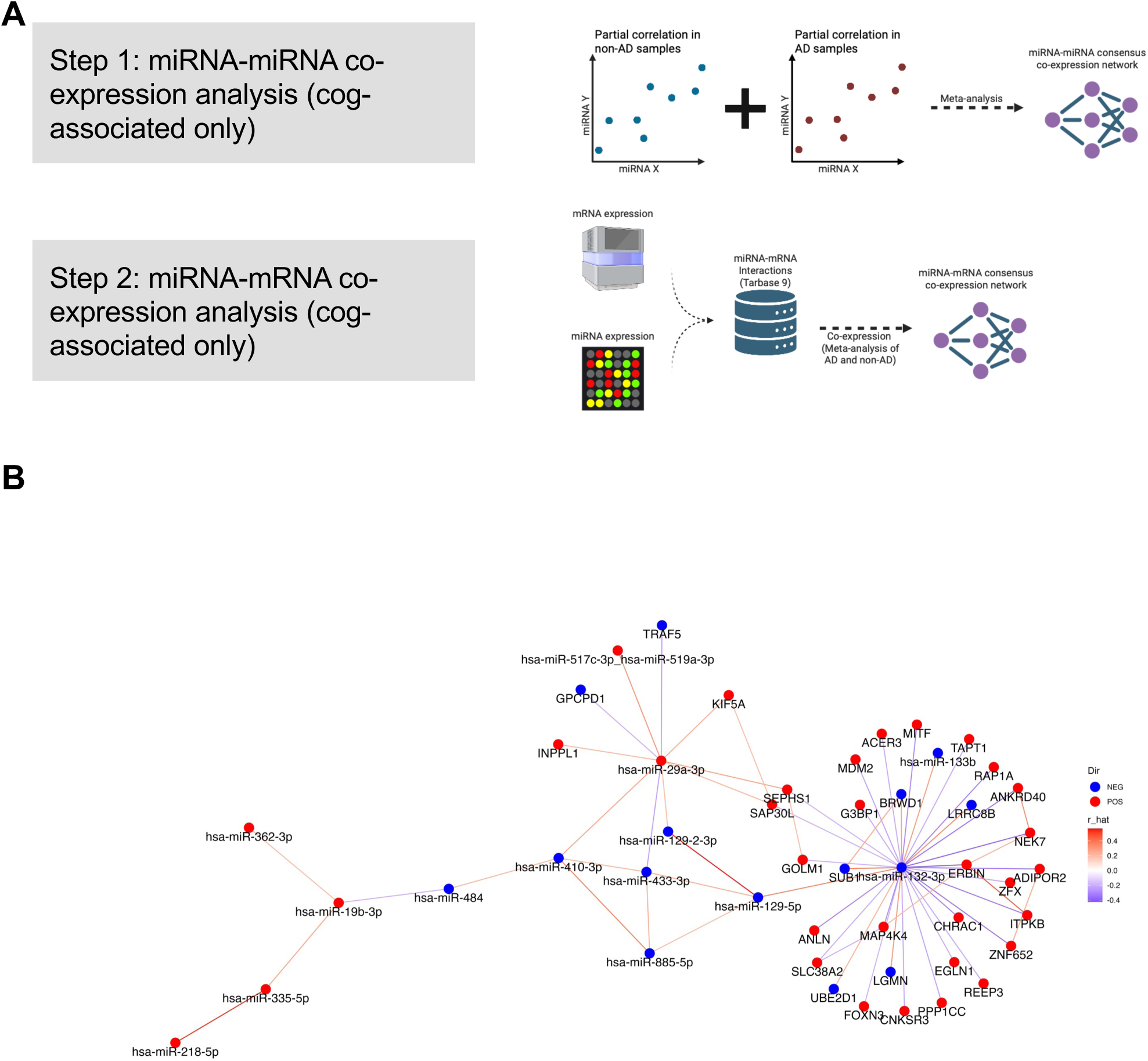
A cognitive-associated miRNA-mRNA co-expression network from the ROSMAP Cohort. **A.** Schematic for deriving the co-expression network from three different components: 1-miRNA-miRNA co-expression; 2-miRNA-mRNA co-expression; and 3-mRNA-mRNA co-expression. In all components, co-expression was calculated separately for AD and nonAD samples, and then merged into a consensus value using Stouffer’s method. miRNA-miRNA co-expression was derived using the partial correlation analysis of cognitive-associated miRNAs. miRNA-mRNA co-expression was filtered to include interactions between cognitive-associated genes and cognitive-associated miRNAs. mRNA-mRNA interactions were limited to cognitive associated genes that had at least one co-expressed cognitive miRNA. **B.** Co-expression network of cognitive associated miRNAs and targeted mRNAs. Links represent statistically significant co-expression events with a consensus adjusted p-value < 0.05 and absolute correlation value of > 0.20. Condition-specific p-values were calculated from z-scores derived using respective correlation values and degrees of freedom. Consensus p-value were derived by aggregating conditions-specific p-values using Stouffer’s method.

miR-132-3p appeared as a central miRNA in the network, with multiple candidate miRNA–mRNA regulatory events evidenced by co-expression, including ERBB2 interacting protein (ERBIN) and microphthalmia-associated transcription factor (MITF) (**Figure 3B**). We found additional co-expression evidence between the genes that were targeted by miR-132-3p, including the co-activity of ERBIN with kinase-encoding genes *MAP4K4, ITPKB,* and *NEK7*. Additional high-potential targets were identified for miR-29a-3p, including glycerophosphocholine phosphodiesterase 1 (GPCPD1). We also found strong evidence of co-expression of cognitive-associated miRNAs, including a subnetwork with several mutually exclusive correlation events involving miR-410-3p, miR-433-3p, miR-885-5p, miR-129-5p, and miR-29a-3p.

### Functional characterization and assignment of miRNAs

The functional roles of miRNAs are better understood in the context of their associated pathways rather than in terms of their single gene targets. To probe functional roles of resilience- and cognitive-associated miRNAs, we used two systematic approaches: (i) data-driven analysis of co-expression of miRNAs and known pathways and (ii) knowledge-driven analysis of enrichment of known miRNA targets in AD and resilience-associated pathways. Data-driven approaches used the previously described computational method to generate pathway activity profiles from ROSMAP gene expression data mapped to canonical functions annotated in the Molecular Signatures Database (MSigDB,v. 2024) (Liberzon et al., 2011). Pathway co-activity analysis was stratified by AD and non-AD subjects (as described above) followed by consensus meta-analysis of miRNA–pathway co-expression across all subjects.

miR-132-3p had the highest number of correlated pathways (adjusted *p* <0.05) in non-AD subjects and across all subjects (**Figure 4**). Top pathways positively correlated with miR-132-3p included neuronal *ligand-receptor interactions, mitogen-activated protein kinase (MAPK) and MAPK14 (p38-alpha),* and *nitric oxide metabolism.* Top pathways negatively correlated with miR-132-3p included *Rho GTPase, sphingolipid processing,* and *stem cell*. In contrast, miR-362-3p had the greatest number of correlated pathways in AD subjects with only a few correlated pathways in non-AD subjects, suggesting that miRNA–pathway co-activity may depend on disease status. Top pathways positively correlated with miR-362-3p included *tumor necrosis factor* (*TNF) and Toll-like receptors,* and *nuclear factor-ĸB (NFKB) activation*. Pathways negatively correlated with miR-362-3p included *anaphase-promoting complex*/*cell division cycle 20* (*APC/CDC20), urea cycle,* and *protein metabolism*. We used the top 30 pathways (when applicable) to define functional correlates of each miRNA of interest including cognitive-associated, plaque-associated, and sex-specific miRNAs (marked by **\*\*\*** in **Figure 2A**). Sex-specific miR-7-5p correlated positively with the Reelin signaling pathway, which we have previously shown to associate with resilient inhibitory neuronal populations (Mathys et al., 2023; Castanho et al., 2025). Plaque-specific miR-199a-5p correlated with only four pathways including the p38 gamma and delta signaling pathway, a key protein phosphorylation pathway mediating microglial activation. **Table 3** summarizes the top 30 pathways for resilience-associated miRNAs, stratified by AD and non-AD samples. **Supplementary Table S5** comprehensively lists miRNA–pathway correlation assessments and top associated pathways for each miRNA.

**Figure 4.**
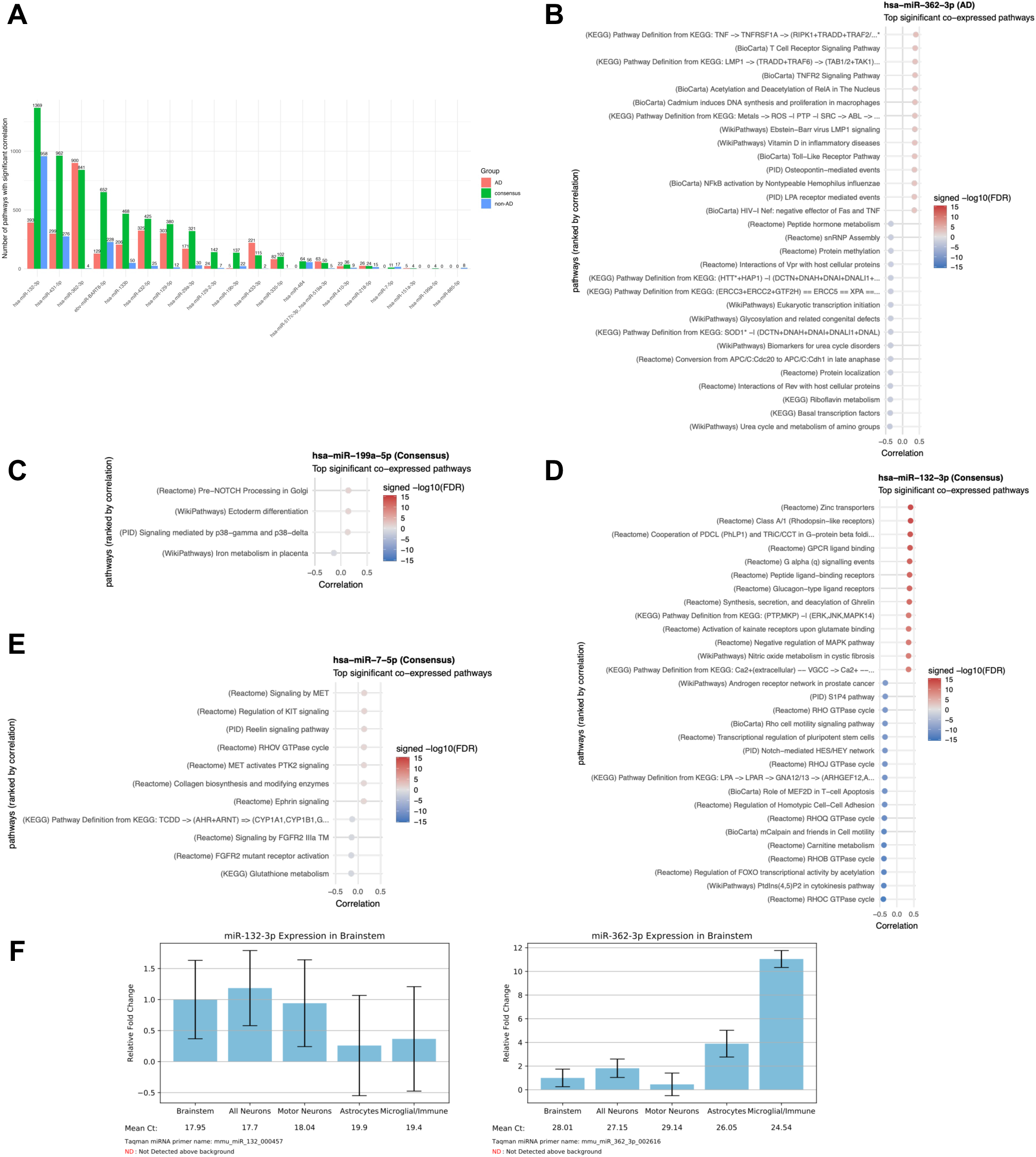
Integrative Co-activity analysis of miRNAs and pathways. **A.** Number of correlated pathways with each AD/resilience associated miRNA based on co-expression analysis of the ROSMAP miRNA and mRNA expression data. Pathway profiles were generated by collapsing gene expression into pathways using the PanomiR and IPAA methods. Correlation coefficients were calculated separately for AD and non-AD samples and evaluated using t-statistics. The values were then merged into a consensus correlation using a weighted Fisher’s transformation and p-values were calculated using Z-scores. **(B-E)** Individual plots representing top 30 significant pathways for the respective miRNAs across the selected conditions. Plots with less than 30 pathways represents miRNAs with less than 30 correlated pathways. **(F)** Cell-specific expression of miR-132-3p and miR-362-3p in mouse brain stem, queried from Hoye et al. data (miRNA.wustl.edu)

To further characterize resilience- and AD-associated miRNAs, we leveraged a database of CNS miRNA profiles (<u>miRNA.wustl.edu</u>) to determine cell-specific restriction of miRNAs based on expression in mice **(Figure 4**, **Table 2, Supplementary Figures)**. miRNAs associated with resilience and cognitive loss seem to be enriched across different neuronal and glial populations: miR-132-3p, miR-129-5p, miR-29a-3p, and miR-218-5p were associated with excitatory neurons, while miR-433-3p and miR-133b were associated with inhibitory neurons.

**Table 2.** Alzheimer’s disease (AD)- and resilience-associated microRNAs (miRNAs). Columns [1] and [2] denote cell types annotated in the CNS microRNA Profiles database, sourced from Hoye et al. 2017 and He et al. 2012, respectively.

| miRNA | AD vs Res | Plaques | Tangles | Cognitive loss | Cell Type [1] | Neuronal Type [2] |
| --- | --- | --- | --- | --- | --- | --- |
| hsa-miR-132-3p | Yes | Yes | Yes | Down | Neurons | Excitatory glutamatergic pyramidal neurons and SOM interneurons |
| hsa-miR-129-5p | Yes | Yes | Yes | Down | Neurons | Excitatory glutamatergic pyramidal neurons (highest) and intermediate in inhibitory neurons |
| hsa-miR-433-3p | Yes |  |  | Down | Neurons | GABAergic and SOM inhibitory neurons (highest) and intermediate in excitatory glutamatergic pyramidal neurons |
| hsa-miR-129-2-3p | Yes | Yes |  | Down | Neurons | Excitatory glutamatergic pyramidal neurons and GABAergic inhibitory neurons |
| hsa-miR-133b |  |  |  | Down | Neurons | GABAergic inhibitory neurons |
| hsa-miR-432-5p | Yes |  |  | Down | Unknown |  |
| hsa-miR-885-5p | Yes |  |  | Down | Astrocytes and microglia |  |
| ebv-miR-BART8-5p |  |  |  | Down | Unknown |  |
| hsa-miR-29a-3p | Yes |  |  | Up | Neurons | Excitatory glutamatergic pyramidal neurons (highest) and intermediate in inhibitory neurons |
| hsa-miR-517c-3p_hsa-miR-519a-3p |  |  |  | Up | Unknown |  |
| hsa-miR-335-5p |  |  |  | Up | Neurons | Excitatory glutamatergic pyramidal neurons, GABAergic and SOM inhibitory neurons |
| hsa-miR-362-3p | Yes |  |  | Up | Microglia and astrocytes, intermediate in neurons |  |
| hsa-miR-19b-3p |  |  |  | Up | Astrocytes |  |
| hsa-miR-484 |  |  |  | Down | Astrocytes and microglia |  |
| hsa-miR-410-3p |  |  |  | Down | Neurons | Pan-neuronal; excitatory glutamatergic pyramidal neurons and inhibitory GABAergic and SOM neurons |
| hsa-miR-218-5p | Yes |  |  | Up | Neurons | Excitatory glutamatergic pyramidal neurons and GABAergic fast-spiking inhibitory neurons |
| hsa-miR-431-5p |  |  |  | Down | Neurons | Pan-neuronal; excitatory glutamatergic pyramidal neurons and inhibitory GABAergic and SOM neurons |
| hsa-miR-7-5p | Male only |  |  |  | Astrocytes | Highly expressed in somatostatin neurons |
| hsa-miR-151a-3p | AD versus control only |  |  |  | Astrocytes |  |
| hsa-miR-199a-5p |  | Yes |  |  | Neurons | GABAergic and SOM inhibitory neurons |

Abbreviations: AD, Alzheimer's disease; GABA, gamma amino butyric acid; Res, resilient; SOM, somatostatin

To systematically characterize mRNAs targeted by resilience miRNAs, we used a knowledge-driven approach in which we investigated miRNAs with enriched targets among pathways associated with loss of cognition (**Figure 5**). Cognitive-associated pathways were identified using a proportional odds model, as described in Methods and previously (Castanho et al., 2025). We used the PanomiR package, which leverages co-expression of pathways to group cognitive-associated pathways into six functionally coherent clusters (Naderi Yeganeh et al., 2023): (1) upregulation of Rho GTPase, matrisome, transforming growth factor β (TGF-β), and yes-associated protein 1 (YAP1) signaling; (2) downregulation of RNA/DNA biosynthesis and metabolism; (3) downregulation mitochondrial processes (4) upregulation of Toll-like receptor (TLR), SMAD signalling, and cytokine activity; (5) dysregulation of cytokine activity and apoptosis; and (6) Upregulation of FoxO signaling and E2F transcription factor network (**Supplementary Figure 2, Supplementary Table S6**).

**Figure 5.**
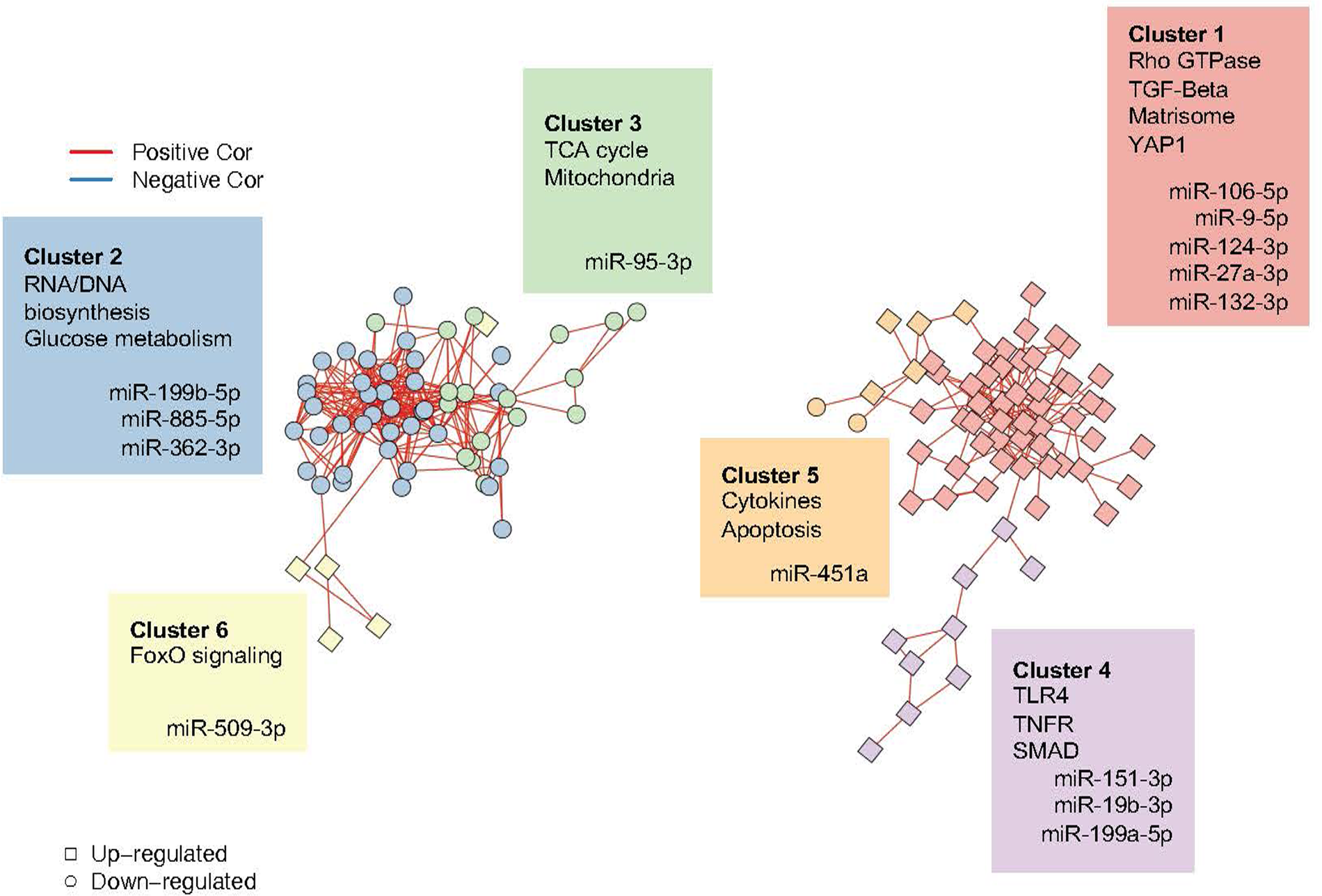
Network analysis of cognitive-associated pathways and targeting miRNAs. **A.** Nodes represent cognitive-associated pathways. Shapes denote their association with the progression of cognitive decline. Links represent statistically significant co-expression of pathways across multiple tissues, derived from the PCxN network. Node colors represent unsupervised clusters generated using the Louvain algorithm. miRNAs associated with each cluster were determined using the PanomiR method. Details on cluster members and specific miRNA targeting events can be found in supplementary figures 2-5.

Leveraging known miRNA–mRNA interactions, we found miRNAs with enrichment of targets across each cluster of pathways, including those with differential expression. miR-132-3p had enriched targets in cluster 1. miR-199b-5p, miR-885, and miR-362 targeted cluster 2. miR-95-3p targeted cluster 3. miR-19b-3p and miR-151-3p targeted cluster 4., and miR-451a targeted cluster 5.

## Discussion

In this study, we systematically mapped the miRNA expression landscape of AD resilience in the human cortex. We identified a distinct set of non-coding regulators that may contribute to maintaining cognitive function despite the presence of AD pathology. By integrating differential gene expression, pathology-specific modeling, and systems-level analyses, we identified novel resilience-associated miRNAs, including miR-362-3p and miR-433-3p. Our analysis also identified resilience association for previously reported AD-associated miRNAs, including miR-132-3p and miR-129-5p. Sex-specific dysregulation of miR-7-5p was observed, with male-specific differential expression in AD versus resilience. These findings indicate that therapeutic strategies aimed at supporting resilience will likely benefit from sex-specific study designs. We also found that miR-199a-5p is exclusively dysregulated in association with plaque pathology but not in association with tau or cognitive loss. Systems biology analysis showed how distinct miRNAs regulate groups of pathways related to matrisome signaling, Rho GTPase family, and cytokines, as well as downregulation of RNA/DNA biogenesis and mitochondrial function.

Despite profound pathological differences between resilient and control subjects, molecular differences were undetectable with our miRNA analysis. These results mirror our prior findings for protein-coding genes (Castanho et al., 2025), supporting the concept that resilience is the ability to maintain a prolonged intermediate state between healthy status and cognitive impairment leading to dementia. In fact, most molecular differences between disease categories were identified when comparing resilience and AD, suggesting that significant molecular changes primarily manifest during widespread neuronal death that accompanies the late-stage loss of cognition. The lack of expression differences between resilient and control subjects suggests that AD pathology in resilient individuals is biologically inert, possibly partially through miRNA-dependent suppression of pathological response pathways. Besides finding new resilience-associated roles for some known AD miRNAs such as miR-132-3p and miR-129-5p, we discovered other miRNAs that are largely uncharacterized in the context of AD. Notably, multiple resilience-associated miRNAs, including miR-362-3p, miR-218-5p, miR-29a-3p, miR-433-3p, and miR-885-5p, showed differential expression only between AD and resilient groups, not between AD and control groups.

miR-132-3p emerged as a central hub within cognitive-associated gene networks. It showed consistent functional correlations in AD, non-AD, and consensus analyses and is distinguished from other resilience miRNAs by the strong cognitive relevance of its predicted targets. miR-132-3p forms a tightly connected co-activity subnetwork with multiple cognition-associated genes with experimentally validated binding sites, including *NEK7, ITPKB,* and *MAP4K4*. These genes also exhibited direct co-expression independent of other network effects, as demonstrated by partial correlation analysis.

miR-362-3p showed AD-specific activation and pathway co-expression compared with subjects without AD diagnosis. In contrast to neuronal miR-132-3p, which strongly correlates with pathways active in both AD and non-AD samples, miR-362-3p showed limited correlation with pathways active in non-AD samples, whereas it has the greatest number of co-activated pathways in AD samples among all resilience-associated miRNAs. miR-362-3p is upregulated in AD compared to resilience and its expression increases along the path of cognitive decline. Querying a cell-specific miRNA expression atlas in mice, miR-362-3p showed restricted expression, mostly in microglia and partly in astrocytes (**Table 2**). Our systems biology analysis suggests that this miRNA is co-activated with, and could regulate, proinflammatory pathways such as TLR and TNF receptor. TNF receptors have been shown to correlate with phosphorylated tau across the AD spectrum and have been suggested as predictors for MCI-to-AD conversion (Zhao et al., 2020). The roles of miR-362-3p in the CNS, especially in the context of AD, are largely unknown. One study showed that miR-362-3p overexpression promotes functional restoration in rat spinal cord injury models through suppression of p38 and ERK pathways (Hu et al., 2019). Taken together, our results point to miR-362-3p activation in microglia during the late stages of AD, highlighting an emerging target for further study of noncoding regulation of microglia in AD. Further studies are needed to determine whether the dysregulation of miR-362-3p and its associated pathways are pathogenic or compensatory in AD.

We identified a sex-specific resilience candidate, miR-7-5p, which demonstrated male-specific dysregulation in AD versus resilience. miR-7-5p also showed a suggestive level of difference between males and females across AD conditions. To our knowledge, this is the first report of a sex-specific miRNA associated with resilience in AD. Interestingly, previous research from our group on sex-dependent miRNA expression following stress identified rno-miR-7b (a mostly conserved homologue of miR-7-5p) as being dysregulated by adolescent social isolation stress in male rats, but not in females (Mavrikaki et al., 2019). We also found multiple other miRNAs with suggestive evidence for sex-specific dysregulation. miR-7 is an evolutionarily conserved miRNA expressed across several brain regions, most prominently in the hypothalamus, and interacts with the neuroendocrine system (Zacharjasz et al., 2024).

Study of sex-specific AD pathways is an emerging domain that could improve understanding of the disease and advance therapeutic approaches. A recent review and critical meta-analysis found minimal reported sex-based differences for miRNAs in AD despite extensive evidence on sex-specific regulation (Llera-Oyola et al., 2024). The common practice of adjusting for sex as a covariate in statistical models obscures sex-specific transcripts by focusing on shared miRNA signatures common in males and females. This is notable given the evidence that AD involves broad sex-specific regulatory programs (Wan et al., 2020). Most of our significant sex-based differences were detected when comparing male and female AD subjects. We did not find any sex-specific differences between males and females in either resilience or control groups. This suggests that sex-specific programs are initiated or become more pronounced during later stages of AD. Our findings demonstrate that both sex-independent and sex-dependent mechanisms contribute to resilience, highlighting the necessity of sex-stratified analyses to reveal sex-specific pathways underlying resilience to AD. Our candidate sex-specific miRNAs, including miR-218-5p and miR-7-5p, may prove to be valuable targets for discovering sex-based regulation of resilience to AD.

Integrative network analysis of miRNAs and gene expression showed persistent co-regulation among resilience miRNAs. In particular, miR-132-3p appeared as a central regulator of cognitive-associated genes in our study, concordant with previous reports on the roles of this miRNA (Salta et al., 2016). miR-132-3p has been shown to interact with BACE1 regulation of APP cleavage and sphingosine kinase, and it was reported as a key miRNA in neuroprotection and hippocampal neurogenesis (Salta et al., 2016; El Fatimy et al., 2018; Hadar et al., 2018; Lauretti et al., 2021; Walgrave et al., 2021). Here, we showed that miR-132-3p is closely associated with the resilience process and is also associated with major pathological and clinical aspects of AD: plaques, tau, and cognitive decline. In addition, miR-129-5p closely follows miR-132-3p patterns in its association with AD progression, evidenced by a significant correlation. Our study showed a close interaction between miR-132-3p and miR-129-5p. Other independent analyses of ROSMAP cohort data have repeatedly found miR-132-3p and miR-129-5p to be associated with the slope of cognitive trajectory across longitudinal visits and across disease stages, providing additional evidence for a resilience function of these miRNAs (Wingo et al., 2022; Han et al., 2024). Another recent study showed that miR-129-5p and miR-132-3p are differentially expressed across the entorhinal cortex and superior temporal gyrus in AD (Dobricic et al., 2022). Additional evidence points to co-dysregulation of these two miRNAs across neurodegenerative diseases (Luo et al., 2025). Our functional analysis demonstrated that both miR-129-5p and miR-132-3p strongly associate with activation of zinc transporters and suppression of Rho GTPase and sphingosine-1 phosphatase (S1P4) pathways. These two miRNAs appear to connect the components of AD pathogenesis and resilience, as they strongly co-activate and may be co-regulated to drive AD progression. Together, our results place miR-129-5p and miR-132-3p as interconnect pair of central resilience factors with high potential as therapeutic targets.

Our analysis isolated miR-199a-5p as a miRNA with a plaque-specific association, potentially showing that the Aβ-linked expression of this miRNA is independent from cognitive decline and tau. Although the exact role of miR-199a in plaque pathology remains unclear, it is involved in blood-brain barrier integrity and response to cerebral ischemic injury (Ni et al., 2024). Recent reports also show that miR-199a-5p interacts with *GSK3B, SIRT1,* and *BDNF.* Although these interactions were identified in non-AD experimental systems, each of these targets has independently been linked to AD progression and resilience (Chen et al., 2018; Liu et al., 2021; Yang et al., 2023). Taken together, the evidence suggests that miR-199a-5p may confer protective roles in AD by directly interacting with Aβ pathology.

We leveraged systems biology methods to assign functions and groups of pathways to resilience-associated miRNAs. miRNAs are inherently capable of targeting several genes across biological functions through non-specific binding (7–8 nucleotide base pairing) in the 3’ UTR of mRNAs. AD-related studies of miRNAs often investigate one or few specific targets for mechanistic characterization, which is contrary to the multi-target nature of miRNA action. Pathway co-activity and targeting analysis of resilience-associated miRNAs revealed convergence on key biological programs regulating neuronal structure, function, and disease response. Multiple miRNAs (miR-29a-3p, miR-129-5p, miR-218-5p, miR-362-3p, and miR-433-3p) correlated with pathways related to immune and inflammatory signaling networks, including TNF, NF-κB, interleukin 12 (IL-12)/signal transducer and activator of transcription factor 4 (STAT4), IL-3, chemokines, TLR, and T-cell receptor signaling pathways, highlighting potential interplay of miRNAs with immune activation in AD (**Figure 4-6**, **Table 3, Table S5, Supplementary Figures**). Cytoskeletal remodeling and synaptic structural pathways, particularly Rho GTPase signaling, extracellular matrix organization, and other matrisome pathways, were correlated with miR-129-5p, miR-132-3p, miR-432-5p, miR-129-2-3p and miR-218-5p in AD, which was also corroborated by multi-pathway enrichment analysis (**Figures 5 and 6**). Additional notable pathways included longevity and stress-resistance programs centered on forkhead box protein O (FOXO) regulation (co-expressed with miR-29a-3p and miR-132-3p), and mitochondrial and metabolic pathways, including nicotinamide adenine dinucleotide (NAD) metabolism, peroxisome proliferator-activated receptor α (PPAR-α) signaling, and fatty acid oxidation (co-expressed with miR-432-5p and miR-885-5p). Together, these findings indicate that AD resilience-associated miRNAs collectively regulate interconnected networks governing immune activation, synaptic structure, and transmission, proteostasis, mitochondrial metabolism, and growth and survival signaling, highlighting coordinated miRNA regulation of pathways that influence the brain’s response and resilience to AD.

**Figure 6.**
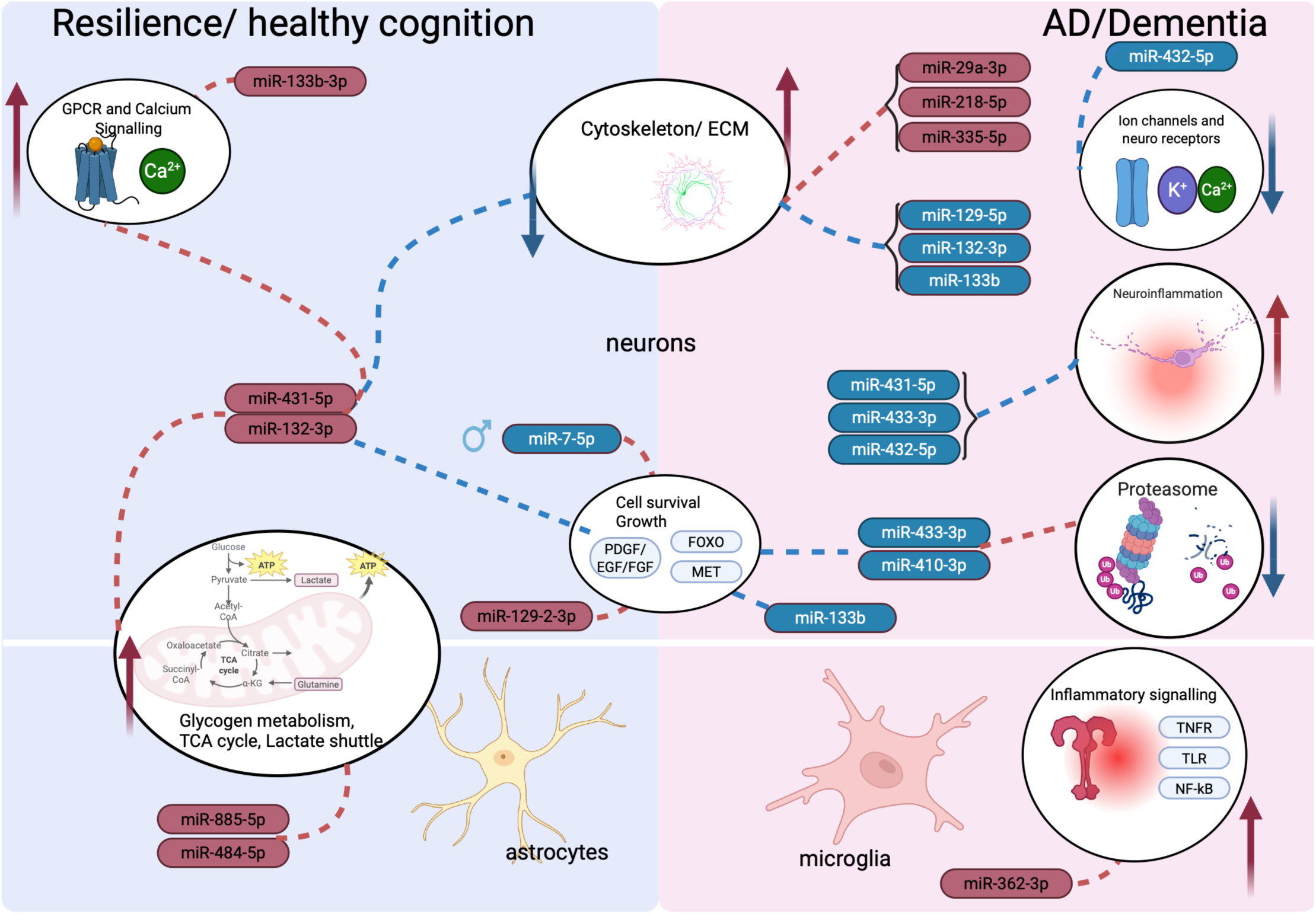
Schematic summary of resilience miRNAs along with their targeted functions, and cell-types.

**Table 3.**
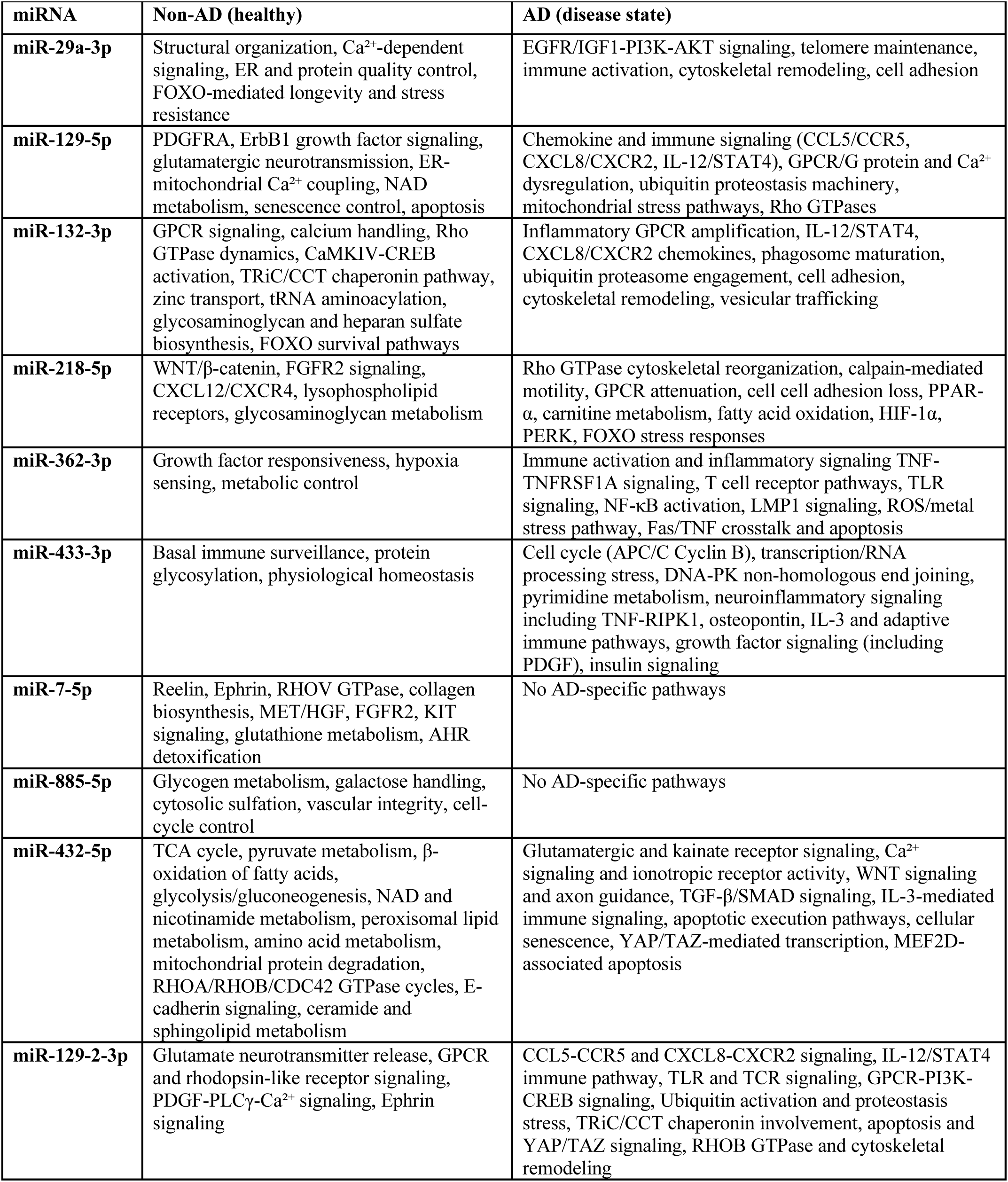
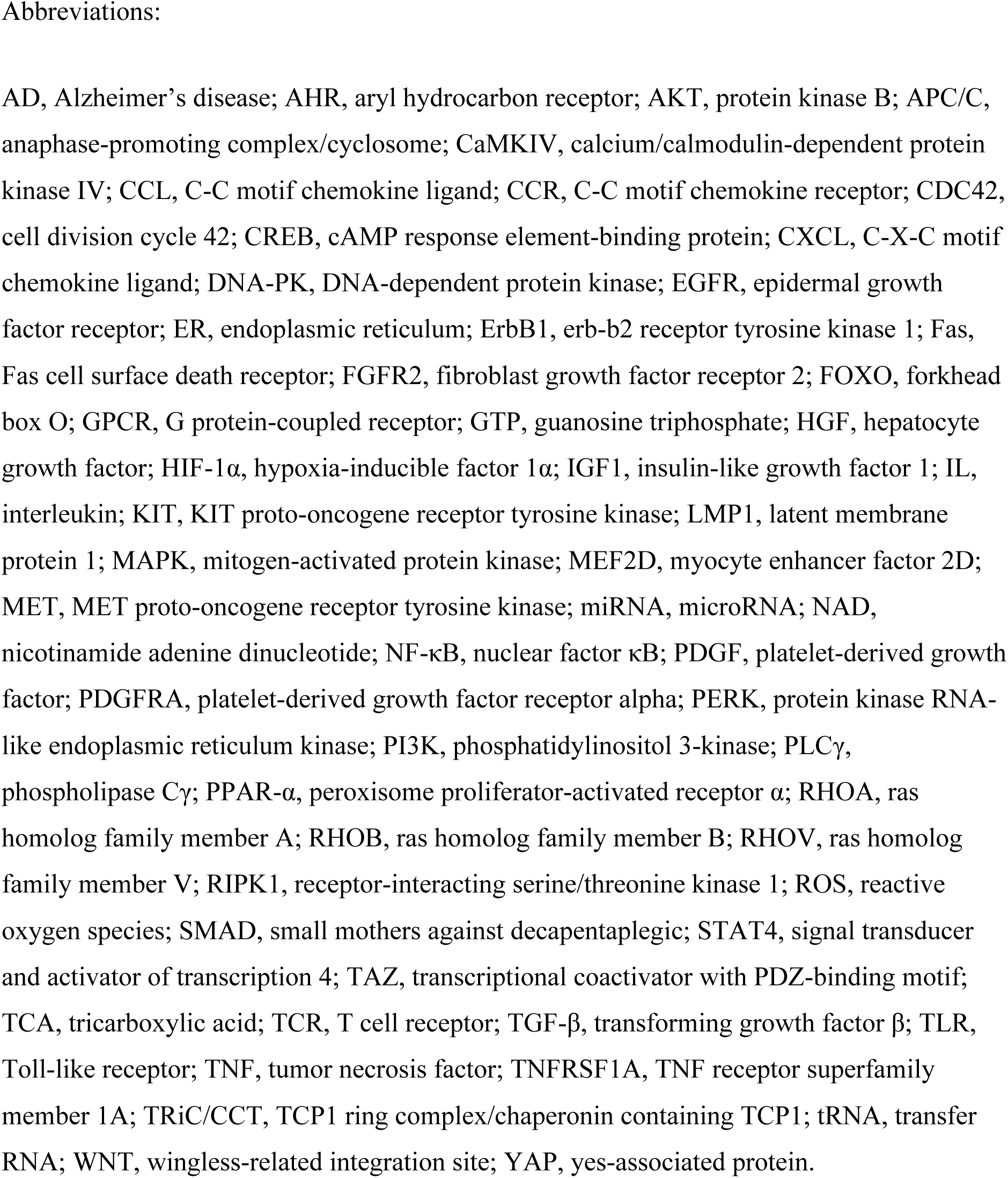
Summary of the top 30 significant pathways associated with resilience microRNAs (miRNAs) [Benjamini-Hochberg false discovery rate (FDR <0.05)]. A full list of pathways is provided in Supplementary Table S5.

| miRNA | Non-AD (healthy) | AD (disease state) |
| --- | --- | --- |
| **miR-29a-3p** | Structural organization, Ca<sup>2+</sup>-dependent signaling, ER and protein quality control, FOXO-mediated longevity and stress resistance | EGFR/IGF1-PI3K-AKT signaling, telomere maintenance, immune activation, cytoskeletal remodeling, cell adhesion |
| **miR-129-5p** | PDGFRA, ErbB1 growth factor signaling, glutamatergic neurotransmission, ER-mitochondrial Ca<sup>2+</sup> coupling, NAD metabolism, senescence control, apoptosis | Chemokine and immune signaling (CCL5/CCR5, CXCL8/CXCR2, IL-12/STAT4), GPCR/G protein and Ca<sup>2+</sup> dysregulation, ubiquitin proteostasis machinery, mitochondrial stress pathways, Rho GTPases |
| **miR-132-3p** | GPCR signaling, calcium handling, Rho GTPase dynamics, CaMKIV-CREB activation, TRiC/CCT chaperonin pathway, zinc transport, tRNA aminoacylation, glycosaminoglycan and heparan sulfate biosynthesis, FOXO survival pathways | Inflammatory GPCR amplification, IL-12/STAT4, CXCL8/CXCR2 chemokines, phagosome maturation, ubiquitin proteasome engagement, cell adhesion, cytoskeletal remodeling, vesicular trafficking |
| **miR-218-5p** | WNT/β-catenin, FGFR2 signaling, CXCL12/CXCR4, lysophospholipid receptors, glycosaminoglycan metabolism | Rho GTPase cytoskeletal reorganization, calpain-mediated motility, GPCR attenuation, cell cell adhesion loss, PPAR-α, carnitine metabolism, fatty acid oxidation, HIF-1α, PERK, FOXO stress responses |
| **miR-362-3p** | Growth factor responsiveness, hypoxia sensing, metabolic control | Immune activation and inflammatory signaling TNF-TNFRSF1A signaling, T cell receptor pathways, TLR signaling, NF-κB activation, LMP1 signaling, ROS/metal stress pathway, Fas/TNF crosstalk and apoptosis |
| **miR-433-3p** | Basal immune surveillance, protein glycosylation, physiological homeostasis | Cell cycle (APC/C Cyclin B), transcription/RNA processing stress, DNA-PK non-homologous end joining, pyrimidine metabolism, neuroinflammatory signaling including TNF-RIPK1, osteopontin, IL-3 and adaptive immune pathways, growth factor signaling (including PDGF), insulin signaling |
| **miR-7-5p** | Reelin, Ephrin, RHOV GTPase, collagen biosynthesis, MET/HGF, FGFR2, KIT signaling, glutathione metabolism, AHR detoxification | No AD-specific pathways |
| **miR-885-5p** | Glycogen metabolism, galactose handling, cytosolic sulfation, vascular integrity, cell-cycle control | No AD-specific pathways |
| **miR-432-5p** | TCA cycle, pyruvate metabolism, β-oxidation of fatty acids, glycolysis/gluconeogenesis, NAD and nicotinamide metabolism, peroxisomal lipid metabolism, amino acid metabolism, mitochondrial protein degradation, RHOA/RHOB/CDC42 GTPase cycles, E-cadherin signaling, ceramide and sphingolipid metabolism | Glutamatergic and kainate receptor signaling, Ca<sup>2+</sup> signaling and ionotropic receptor activity, WNT signaling and axon guidance, TGF-β/SMAD signaling, IL-3-mediated immune signaling, apoptotic execution pathways, cellular senescence, YAP/TAZ-mediated transcription, MEF2D-associated apoptosis |
| **miR-129-2-3p** | Glutamate neurotransmitter release, GPCR and rhodopsin-like receptor signaling, PDGF-PLCγ-Ca<sup>2+</sup> signaling, Ephrin signaling | CCL5-CCR5 and CXCL8-CXCR2 signaling, IL-12/STAT4 immune pathway, TLR and TCR signaling, GPCR-PI3K-CREB signaling, Ubiquitin activation and proteostasis stress, TRiC/CCT chaperonin involvement, apoptosis and YAP/TAZ signaling, RHOB GTPase and cytoskeletal remodeling |

Abbreviations:

The definition of AD resilience is not unified across studies, which may complicate cross-study comparisons. Some recent reports have adopted a continuous definition of resilience (e.g., actual cognitive score versus expected cognitive score), while other have used discrete definitions with pathological cut-offs (Mathys et al., 2023; Castanho et al., 2025). We adopted a stringent discrete approach that measures extreme ends of the pathological spectrum for defining resilience and AD (Castanho et al., 2025). A recent transcriptome-wide study of resilience miRNAs, employing a partially different definition of resilience compared to ours, found involvement of the MAPK pathway in genes targeted by resilience miRNAs in postmortem posterior cingulate cortex (Counts et al., 2025). While this study also showed differential miRNAs across AD, resilient, and MCI subjects, only a few miRNAs overlap with our findings, which is potentially due to differences in definitions, regions, and technical methods used. As a methodological safeguard to ensure the robustness of our findings, we directly modelled cognition across the population without binarizing subjects into resilience, AD, and control categories. Our results across different analytical setups remain consistent and produce a highly overlapping set of miRNAs (**Figure 2A**).

This study extensively leverages co-expression and target enrichment to characterize miRNAs and their interacting genes and pathways (**Figure 6**, **Table 3, Table S5**). However, identifying causal and regulatory relationships will require in-depth experimental designs in model systems that capture the facets explored in this study: pathological progression, neurodegeneration, and human-relevant physiological environments. Additionally, miRNA regulation can be cell- and tissue-specific. Public miRNA–mRNA interaction databases often include canonical relationships that may not reflect interactions occurring in specific cell types or human tissues. Experimental approaches, such as miRNA perturbation experiments and cross-linking immune precipitation sequencing (CLIP), are required to more accurately characterize the roles of the miRNAs discussed here.

Our findings offer a new, compelling set of candidate therapeutic targets for emulating natural resilience to AD. Since the discovery of miRNAs in humans two decades ago (Pasquinelli et al., 2000), they have emerged as key elements in the molecular etiology of neurodegenerative disease. Notable examples of brain miRNAs include miR-132 miR-9, and miR-124, which have been reported in multiple diseases including AD, Parkinson’s disease, and Huntington’s disease (Fukuoka et al., 2018; Han et al., 2019; Walgrave et al., 2021). The translational implications of these findings and the logistics of using miRNAs as therapies are less developed. Emerging examples include small molecules targeting miRNAs [e.g., PrimeC (NeuroSense), trial NCT06185543] and microRNA mimics/inhibitors [e.g., MRG-107 (Viridian Therapeutics)]. miRNAs were also central to a recent breakthrough in Huntington’s disease therapy, evidenced by the promising results from the AMT-130 trial leveraging a synthetic miRNA to silence the *HTT* gene (eClinicalMedicine, 2025). Parallel advances in neurodegenerative disease biology, RNA therapeutics, and RNA delivery methodologies embolden miRNAs as next-generation therapeutic targets in neurodegeneration.

### Conclusion

Our results demonstrate broad association and involvement of miRNAs across AD pathogenesis, functional responses, and cell types (**Figure 6**). Several miRNAs show differential expression linked to resilience to AD, with additional evidence for sex- and pathology-specific regulation. Given the endogenous expression of these miRNAs and their association with resilience, they provide promising candidate biomarkers for identifying resilient individuals and stratifying disease risk. At the same time, resilience-associated miRNAs are emerging as candidate therapeutic targets for strategies aimed at recapitulating natural protection against AD.

## Methods

### miRNA and mRNA data processing

miRNA expression data were generated by the ROSMAP cohort, retrieved through Synapse.org. ROSMAP miRNA expression data were generated using Nanostring nCounter profiling, derived from post-mortem autopsy DLPFC samples (Patrick et al., 2017).

miRNA expression data (*n* = 704 samples, measuring 309 miRNAs) provided and normalized by ROSMAP were used for further downstream analysis, which was filtered for removal of batch effects using the ComBat method (Patrick et al., 2017). miRNA probes and annotations were updated by matching probes with mirBase 22.1 annotation (Kozomara and Griffiths-Jones, 2014). Additional raw and processed expression matrices were retrieved along with associated clinical, demographic, neuropathological, and technical metadata. Samples were filtered based on data availability and quality control metrics, including RNA integrity number (RIN) ³ 5 and post-mortem interval (PMI). The final miRNA expression dataset included 568 subjects.

To analyze mRNA expression, we used RNA-sequencing profiles (*n* = 631) of DLPFC tissues from the ROSMAP cohort, retrieved from the AMP-AD knowledge portal on Synapse.org. Preprocessing, normalization, and subject classification were extensively discussed in our previous publication (Castanho et al., 2025).

### Sample classification

Subjects were categorized into AD, resilient, control, and presymptomatic AD groups based on our previously described criteria integrating cognitive impairment (CI) diagnosis, neuritic plaque burden, Braak tangle stage, and age. Briefly, the control category was characterized by no cognitive impairment (NCI), low plaque pathology [Consortium to Establish a Registry for Alzheimer’s Disease (CERAD) score: sparse/none], and low tangle pathology (Braak stage 0, I, II). The resilient category exhibited NCI, high plaque pathology (CERAD score: frequent/moderate), high tangle pathology (Braak score III–VI), and age at the time of death >80. The AD category exhibited Alzheimer’s dementia with no other conditions contributing to CI, high plaque pathology (CERAD score: frequent/moderate), high tangle pathology (Braak score III–VI), and age at the time of death >80. The presymptomatic category exhibited MCI, high plaque pathology (CERAD score: frequent/moderate), and high tangle pathology (Braak score III–VI). All other subjects not belonging to any of the categories were classified as “other”. The age cut-off for the resilience group represents exposure to aging, which is the strongest risk factor for AD.

### Differential miRNA expression analysis

Linear models, implemented in Limma, were used to perform miRNA differential expression analysis between disease categories, adjusting for covariates (Ritchie et al., 2015). Linear models included additional terms to adjust for sex, RIN, age at the time of death, PMI, and *APOE* genotype. Contrasts tested included AD versus resilient, AD versus control, resilient versus control, presymptomatic comparisons, and AD versus non-AD (control + presymptomatic + resilient). P values for each contrast were derived using a moderated *t*-test, implemented in Limma. P values were then adjusted for multiple hypothesis testing using the Benjamini-Hochberg FDR method. For further downstream analysis, residual gene expression data were generated after adjusting for the following covariates: sex, RIN, age at the time of death, PMI, and *APOE* genotype.

### Ordinal modeling of miRNAs associated with AD progression

Residual miRNA expression was analyzed alongside clinical covariates (i.e., plaques, tangles, and cognitive status). Analyses were restricted to participants with cognitive status codes 1, 2, or 4 (no impairment, MCI, AD dementia). For each miRNA, proportional odds regression was fit using a generalized linear model with a cumulative logit link function (proportional odds assumption) using the Vector Generalised Additive Model (VGAM) R package (Agresti, 2010; Yee, 2020). Three ordinal outcomes were modeled separately as a function of miRNA expression: cognitive diagnosis, neurofibrillary tangle burden, and neuritic plaque burden. For each outcome, we extracted the coefficient for the miRNA term, along its associated p value and R-squared goodness of fit for latent variable models. P values were adjusted for multiple testing using FDR. We also implemented an additional proportional odds model to test cognition (dependent variable) as a function of miRNA expression, adjusted for the levels of plaques and tangles, to evaluate the relationship of miRNAs with cognitive status (AD, MCI, healthy) independent of plaques and tangles. To plot each outcome, expression was normalized within each miRNA by subtracting the mean expression of the lowest pathology/impairment group (e.g., no impairment for cognition, no tangles for Braak). Group means were plotted across ordered categories, and miRNAs were labeled at the most severe category (e.g., AD dementia, Braak VI).

### Sex-specific differential miRNA expression analysis

Sex-specific effects were evaluated using a linear model with miRNA expression (dependent variable) where independent variables included age, PMI, APOE genotype, RIN and an interaction term between diagnosis (e.g., AD, resilience) and sex. Contrasts tested disease effects within each sex, sex differences within diagnostic groups, and sex-by-diagnosis interactions. P values for each contrast were derived using a moderated *t*-test, implemented in the Limma package. Results were compared with outcomes from proportional odds models and differential expression analysis of all subjects.

### Integrated miRNA–mRNA co-expression network analysis

Co-expression analysis paired with known annotated miRNA–mRNA interactions was used to derive a co-activity transcriptomic network in AD and control groups. Residual miRNA expression data were used to construct co-expression analyses stratified by cognitive diagnosis. Samples were partitioned into AD (diagnosis of AD dementia) and non-AD (NCI or MCI), with all analyses restricted to these groups. miRNA expression was z-scaled per miRNA. Three components of the network were generated sequentially: miRNA–miRNA co-activity, miRNA–mRNA co-activity, and mRNA–mRNA co-activity.

#### miRNA–miRNA analysis

Analysis was limited to miRNAs of interest, defined as the union of significant AD versus resilience miRNAs and cognition-associated miRNAs from proportional odds models (FDR <0.05). Partial correlations between all pairs of miRNAs were computed separately in AD and non-AD groups using the inverse covariance (precision) matrix. The significance of each pair was evaluated using *t*-test with n – p degrees of freedom. P values were adjusted using FDR. AD and non-AD edges were joined and summarized with a consensus correlation meta-estimate using a weighted Fisher’s transformation (inverse tangent hyperbolic).

#### miRNA–mRNA analysis

Analysis was limited to miRNAs of interest, defined as the union of significant AD versus resilience miRNAs and cognition-associated miRNAs from proportional odds models (FDR <0.05). Correlation analysis was performed on samples that had concordant miRNA and mRNA expression. We selected miRNA–mRNA pairs of interest for assessment based on interactions annotated in TarBase V9 database, which contains experimentally validated interactions. TarBase pairs were restricted to high-confidence, primary interaction (direct binding), negative effect of expression, and 3′ UTR interactions in *Homo sapiens* with microT score >0.5. For each TarBase miRNA-gene pair, Pearson correlations were computed within AD and non-AD groups; p values were obtained from *t*-tests and adjusted with FDR within group. AD and non-AD results were merged into a consensus meta-estimate as described above. Results were filtered for significant edges and significant cognition-associated genes (FDR <0.01). Cognition-associated genes were generated as described above in the miRNA proportional odds model and as previously reported (Castanho et al., 2025).

#### mRNA–mRNA analysis

mRNA features were restricted to genes found in miRNA–mRNA resilience correlations as described above. Residual mRNA expression was subset to these genes and stratified by AD versus non-AD. Partial correlation between mRNA pairs were generated as described above. AD and non-AD results were merged into a consensus meta-estimate as described above. The components were then merged into a network with edges representing significant consensus correlation (FDR <0.05) with an absolute value >0.2 (|r| >0.2).

### Functional pathway analysis and miRNA regulatory prioritization

Pathway activity scores were generated from ROSMAP gene expression data using the PanomiR package (Naderi Yeganeh et al., 2023, 2025). Pathway scores were generated using the MSigDB2024 dataset (Liberzon et al., 2011). miRNA–pathway co-activity was generated for AD and non-AD samples as described above. Consensus correlation and its significance analysis were performed using meta-analysis estimates as described above.

Pathways associated with cognition were determined using a proportional odds model as described above. Cognitive-associated pathways were clustered using the default parameters of the PanomiR package, which uses Louvain clustering. miRNA–mRNA interactions for PanomiR were resourced from TarBase V9 using the parameters described above (Skoufos et al., 2024). PanomiR results were limited to miRNAs that were expressed in the ROSMAP dataset and annotated in the TarBase background. PanomiR p-values were generated for the top six clusters of cognitive-associated pathways. We reported the top 30 predicted miRNAs for each cluster and defined significant miRNAs as those with FDR <0.05.

### Cell-type annotation of miRNAs

Selected miRNAs were characterized for cell-type specific expression using the CNS microRNA Profiles, a public database for brain miRNA expression in mice accessed via web portal/database (miRNA.wustl.edu), (He et al., 2012; Hoye et al., 2017). The database provides miRNAs across astrocytes, microglia, and neurons in brainstem and spinal cord. It also provides miRNA expression profiles across neuronal subtypes (Gad2, CamkIIa, Pv, Som) in the neocortex.

### Visualization and data analysis

Volcano plots were generated with EnhancedVolcano, with points colored by FDR significance and effect direction and labeled for FDR <0.05. Networks were constructed and visualized using the igraph package (Csardi and Nepusz, 2006). The ComplexHeatmap package was used for heatmap visualization (Gu et al., 2016).

## Statements

### Data availability and Ethics

No animal studies are presented in this manuscript. No human studies are presented in the manuscript. No potentially identifiable images or data are presented in this study. The original contributions presented in the study are included in the article/supplementary material, further inquiries can be directed to the corresponding author/s.

Gene and miRNA expression datasets used in this study are publicly available through the AMP-AD data portal hosted on Synapse under their original ROSMAP accession code syn8456629 under controlled use conditions subject to privacy regulations. The code used for this publication will be shared on GitHub prior to final publication.

### Author contribution

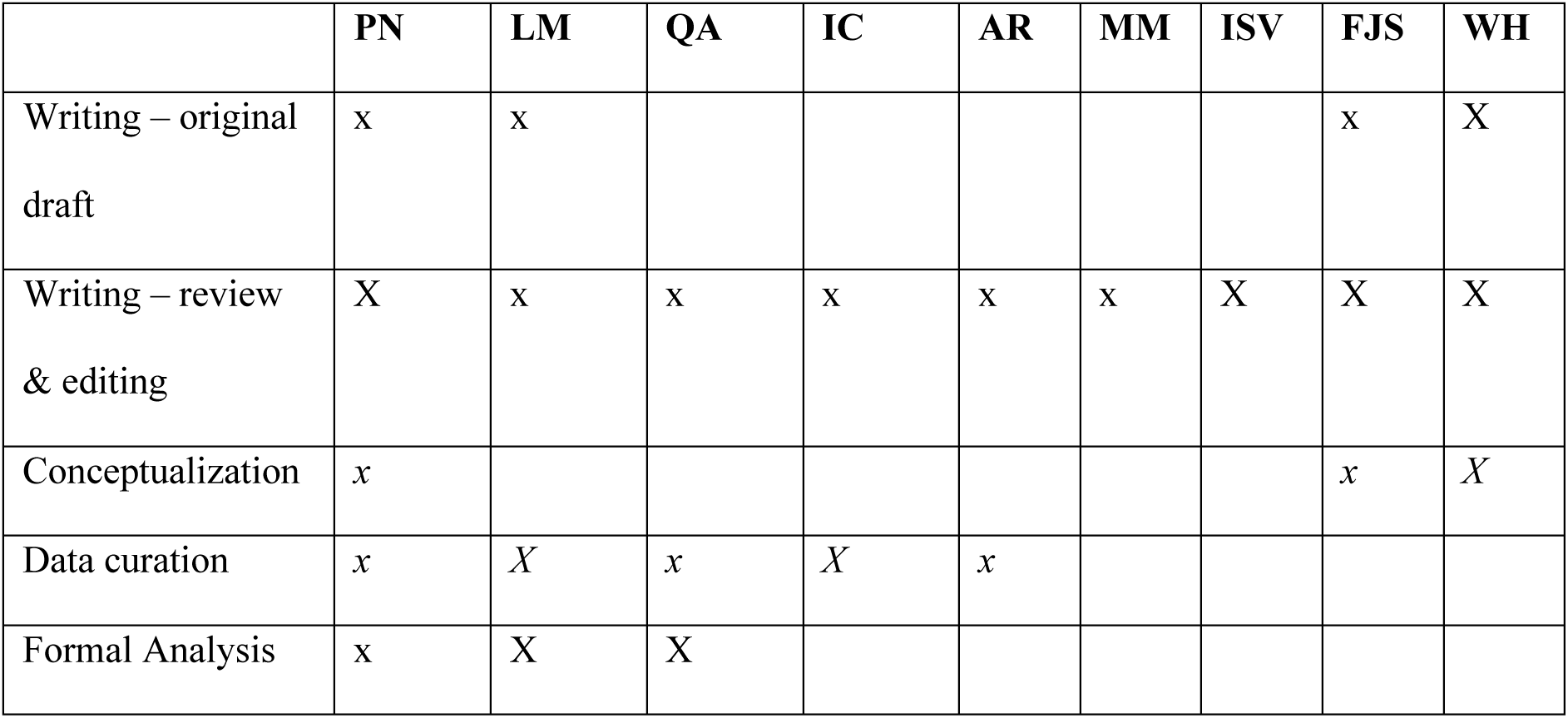
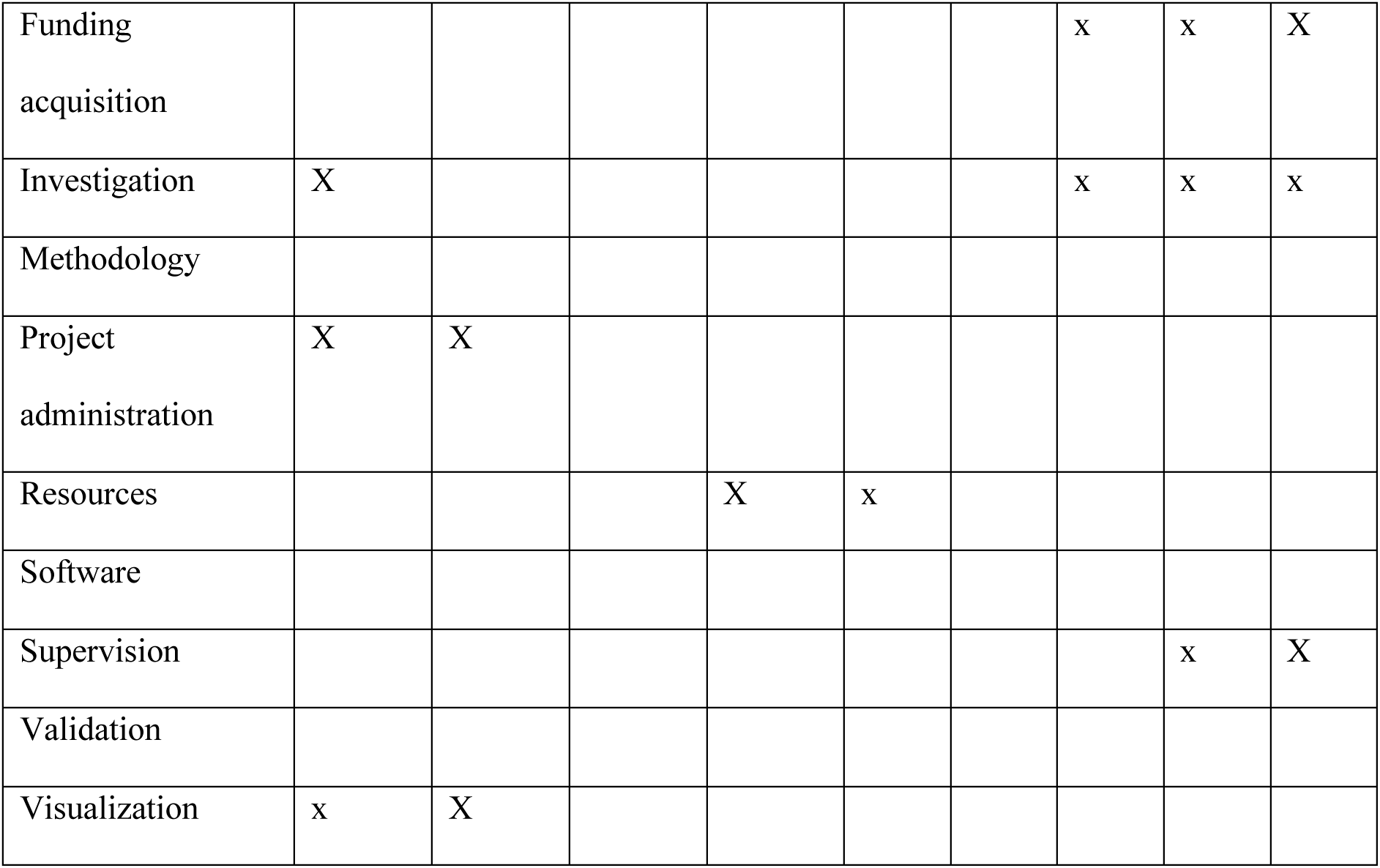

---

Figure 1. Micro RNA (miRNA) expression analysis. **(A)** Study overview. **(B)** Differential expression analysis between Alzheimer’s disease (AD), resilient (Res), and control (Ctr) groups. Differential expression was performed using a linear model accounting for covariates and assessed using a two-sided moderated *t*-test, implemented in the Limma package. Colored points represent significant differential expression [Benjamini-Hochberg false discovery rate (FDR) <0.05]. Dashed lines in each panel represent the p value corresponding to FDR <0.05 cut-off. (DLPFC, dorsolateral prefrontal cortex; mRNA, messenger RNA; ROSMAP, Religious Order Study and Memory and Aging Project) **(C)** Panels show miRNAs with statistically significant trends associated with the progression of cognitive decline and/or AD pathology. Y axes represent the mean of normalized miRNA expression across all samples corresponding to each value on the Y-axis, centered to the lowest category on the Y-axis (e.g., no tangles). Significant trends were determined using a proportional odds model for ordinal categorical regression. The model was implemented with a cumulative logit link using the Vector Generalised Additive Model (VGAM) package. P values were derived from two-sided z-scores of the proportional odds model.

Figure 2. Sex-specific microRNA (miRNA) expression analysis. **(A)** Heatmap showing differential expression p-values across varying contrasts. Sex-specific differential expression analysis was performed using a linear model with an interaction term between covariates for sex and diagnosis, i.e., Alzheimer’s disease (AD), resilient, or control. Each column in the heatmap represents a contrast (All: non-sex specific; Male: males only; Female: females only; M vs F: comparison of males and females). Cell colors represent signed log p values. P values were derived using moderated *t*-test (implemented in the Limma package) and adjusted for multiple comparisons using the Benjamini-Hochberg false discovery rate (FDR) ***: FDR <0.05. ^: FDR <0.2 and *p* <0.05. (**B)** Boxplot of the levels of miRNAs with sex-specific differential expression. P values were derived and adjusted as in **(A)**.

Figure 3. A cognitive-associated microRNA (miRNA)–messenger RNA (mRNA) co-expression network from the Religious Order Study and Memory and Aging Project (ROSMAP) Cohort. **(A)** Schematic for deriving the co-expression network from three different components: 1 – miRNA–miRNA co-expression; 2 – miRNA–mRNA co-expression; and 3 – mRNA–mRNA co-expression. In all components, co-expression was calculated separately for Alzheimer’s disease (AD) and non-AD samples and then merged into a consensus value using Stouffer’s method. miRNA–miRNA co-expression was derived using the partial correlation analysis of cognitive-associated miRNAs. miRNA–mRNA co-expression was filtered to include interactions between cognitive-associated genes and cognitive-associated miRNAs. mRNA–mRNA interactions were limited to cognitive-associated genes that had at least one co-expressed cognitive miRNA. (**B)** Co-expression network of cognitive associated miRNAs and targeted mRNAs. Links represent statistically significant co-expression events with a consensus adjusted p value <0.05 and absolute correlation value of >0.20. Condition-specific p values were calculated from z-scores derived using respective correlation values and degrees of freedom. Consensus p values were derived by aggregating conditions-specific p values using Stouffer’s method.

Figure 4. Integrative co-activity analysis of microRNAs (miRNAs) and pathways. **(A)** Number of correlated pathways with each Alzheimer’s disease (AD)-/resilience-associated miRNA based on co-expression analysis of Religious Order Study and Memory and Aging Project (ROSMAP) miRNA and mRNA expression data. Pathway profiles were generated by collapsing gene expression into pathways using the PanomiR and Integrative Pathway Activity Analysis (IPAA) methods. Correlation coefficients were calculated separately for AD and non-AD samples and evaluated using *t*-statistics. The values were then merged into a consensus correlation using a weighted Fisher’s transformation and p values were calculated using z-scores. **(B**–**E)** Individual plots representing the top 30 significant pathways for the respective miRNAs across the selected conditions. Plots with less than 30 pathways represent miRNAs with less than 30 correlated pathways. **(F)** Cell-specific expression of miR-132-3p and miR-362-3p in mouse brain stem, queried from Hoye et al. data (miRNA.wustl.edu).

Figure 5. Network analysis of cognitive-associated pathways and targeting microRNAs (miRNAs). Nodes represent cognitive-associated pathways. Shapes denote their association with the progression of cognitive decline. Links represent statistically significant co-expression of pathways across multiple tissues, derived from the PCxN network. Node colors represent unsupervised clusters generated using the Louvain algorithm. miRNAs associated with each cluster were determined using the PanomiR method. Details on cluster members and specific miRNA targeting events can be found in supplementary figures. (HDAC, histone deacetylase; MAPK, mitogen-activated protein kinase; Rho GTPase, Ras homolog guanosine triphosphatase; TGF, transforming growth factor; TLR, Toll-like receptor; TNF, tumor necrosis factor; YAP1, yes-associated protein 1)

Figure 6. Schematic summarizing an integrated network of cognition- and resilience-associated microRNAs (miRNAs). Dashed lines denote significant co-expression between miRNAs and pathways. White circles denote functional themes representing pathways. miRNAs overlapping on functions indicate co-expression. Node colors denote up- or down­regulation (red: upregulated, blue: downregulated) in the respective conditions. Pathways converge on key processes including cytoskeletal organization, proteasome, immune signaling, mitochondrial metabolism, and cell survival, emphasizing the intersection of metabolic dysfunction and neuroinflammation in Alzheimer’s disease (AD). Further details on miRNA phenotypes are provided in **Table 2**. (ECM, extracellular matrix; EGF, epidermal growth factor; FGF, fibroblast growth factor; FOXO, forkhead box O; GPCR, G protein-coupled receptor; MET, MET proto-oncogene receptor tyrosine kinase; NF-κB, nuclear factor κB; PDGF, platelet-derived growth factor; TCA cycle, tricarboxylic acid cycle; TLR, Toll-like receptor; TNFR, tumor necrosis factor receptor)

## Supplementary material

- *Supplementary Tables S1-S6*
- *Supplementary Figures S1-S3*

