## Supplementary Figures for "MicroRNA correlates of resilience to Alzheimer’s disease identify candidate therapeutic targets for neuroprotection"

### Supplementary Figure 1

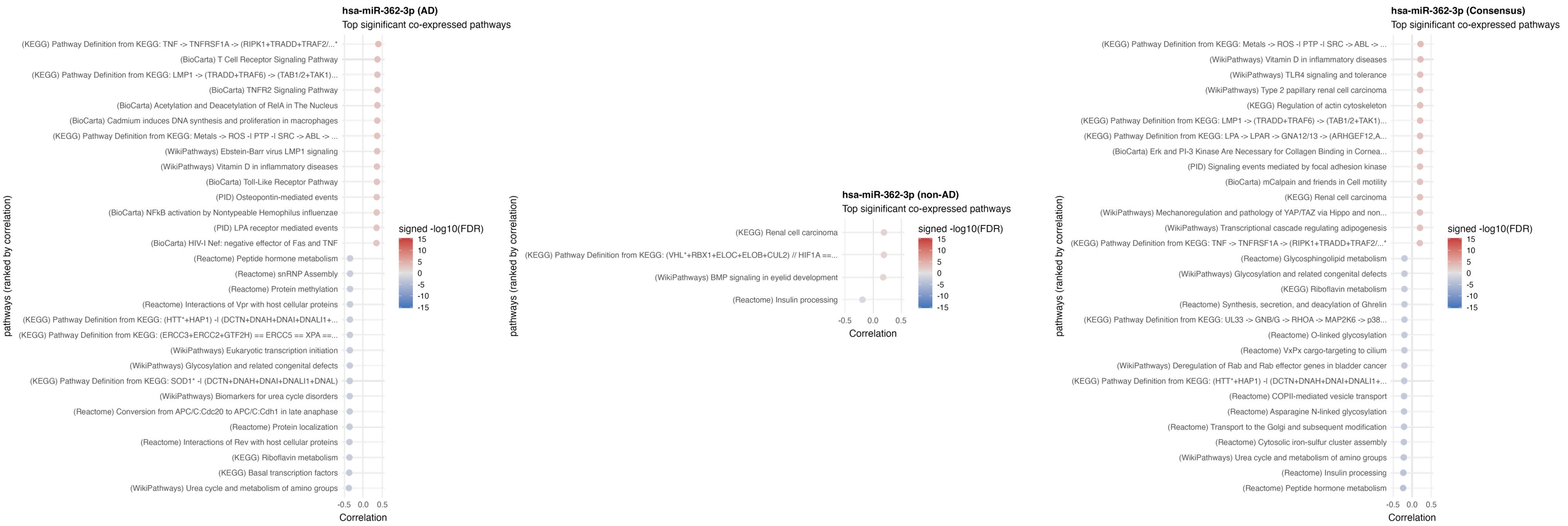


**Supplementary Figure 1. Top correlated pathways with each miRNA in AD and non-AD conditions (related to Figure 4).** The left panel represent AD-specific co-expression. The center panel represents non-AD co-expression. The right panel represents consensus co-expression, derived using meta analysis of AD and non-AD co-expression.

### Supplementary Figure 1 continued

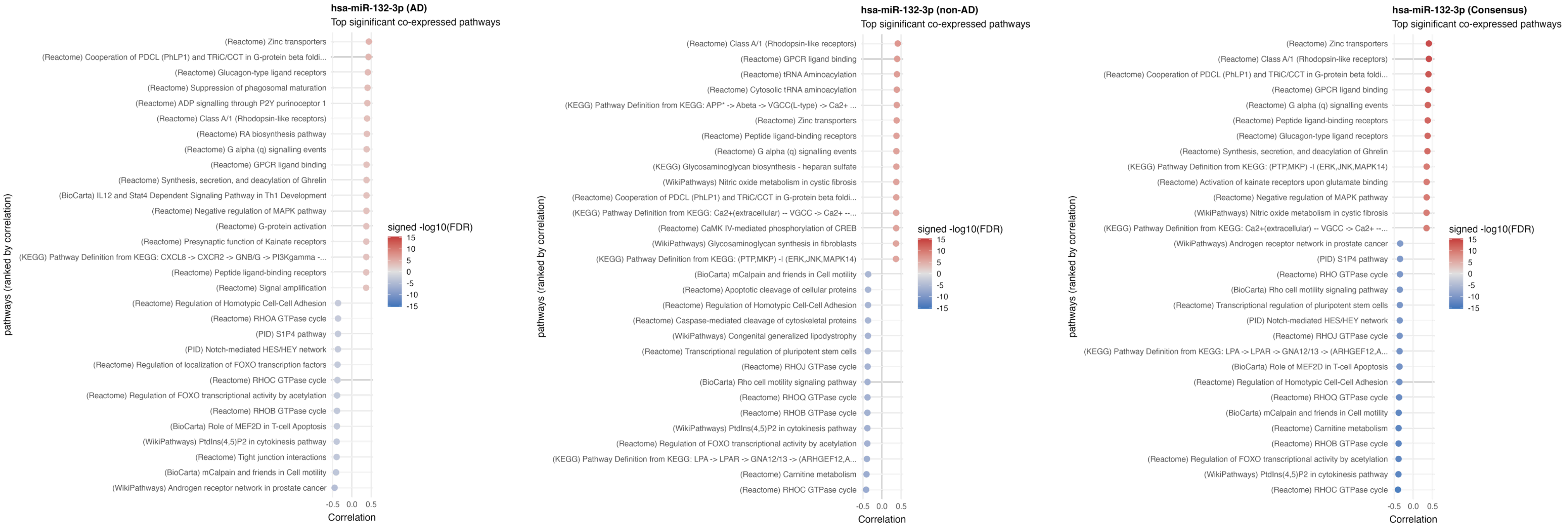

### Supplementary Figure 1 continued

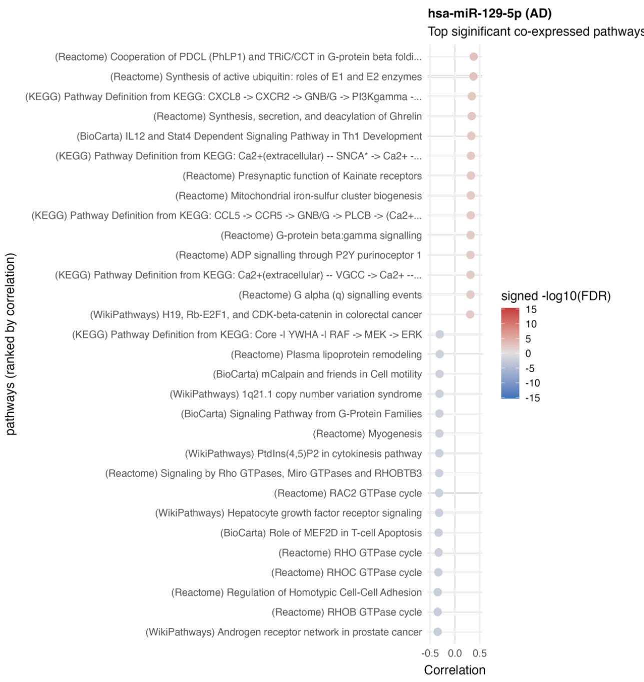


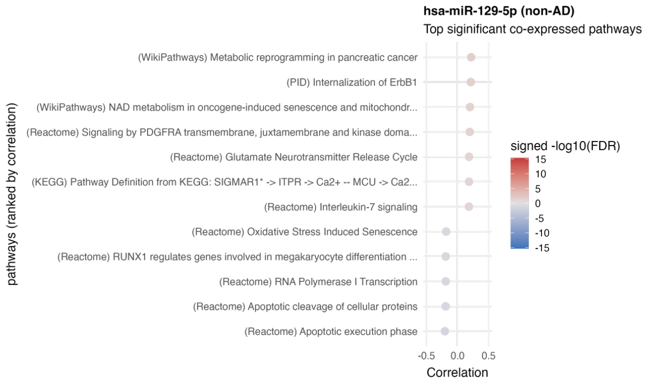


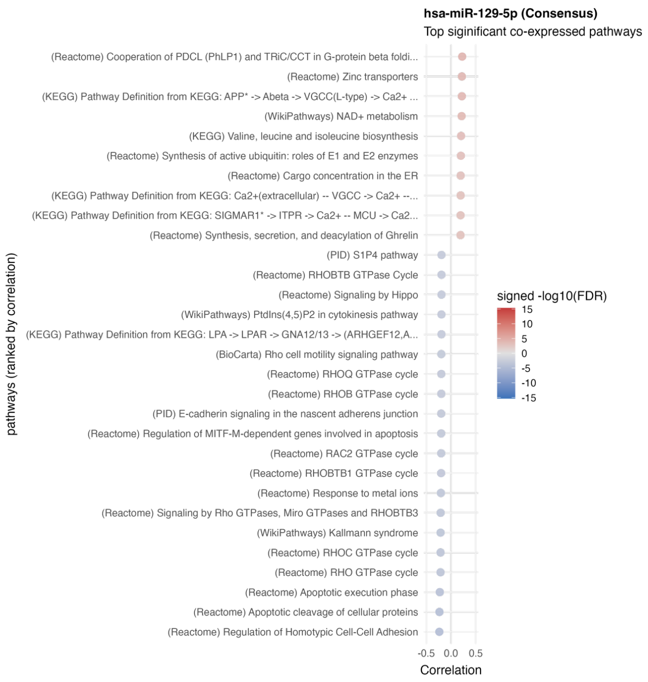

### Supplementary Figure 1 continued

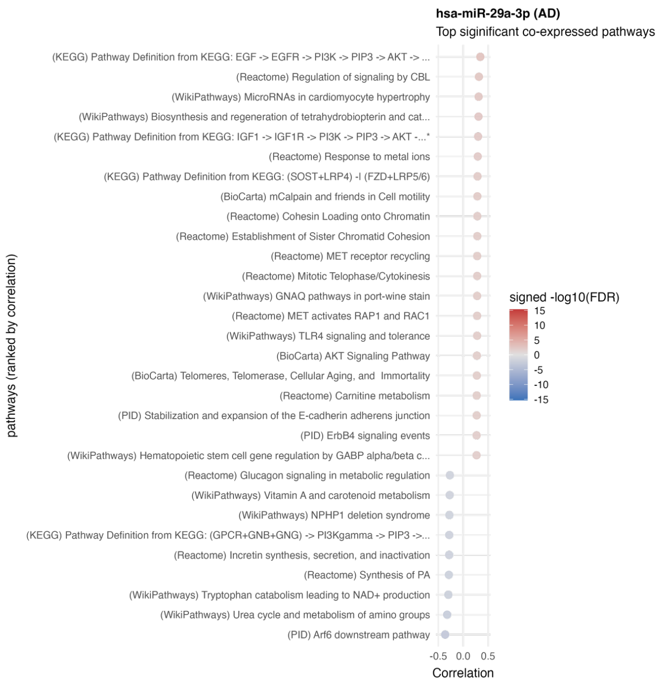


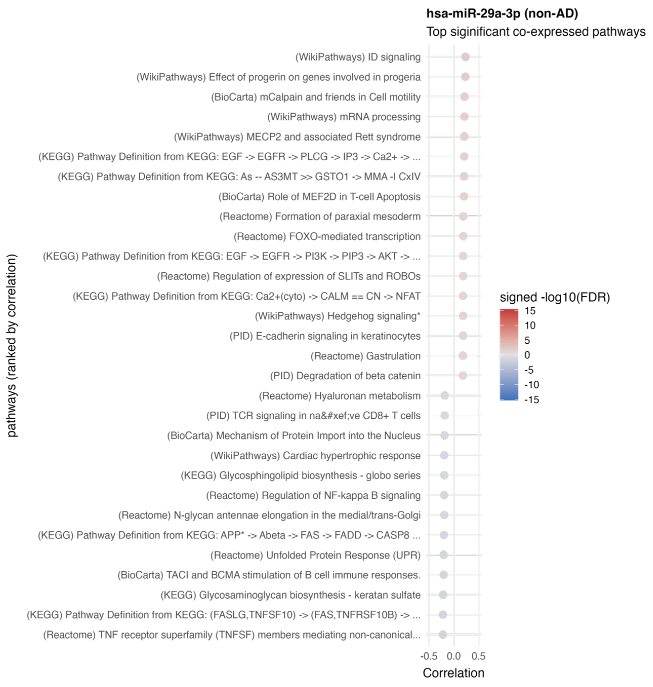


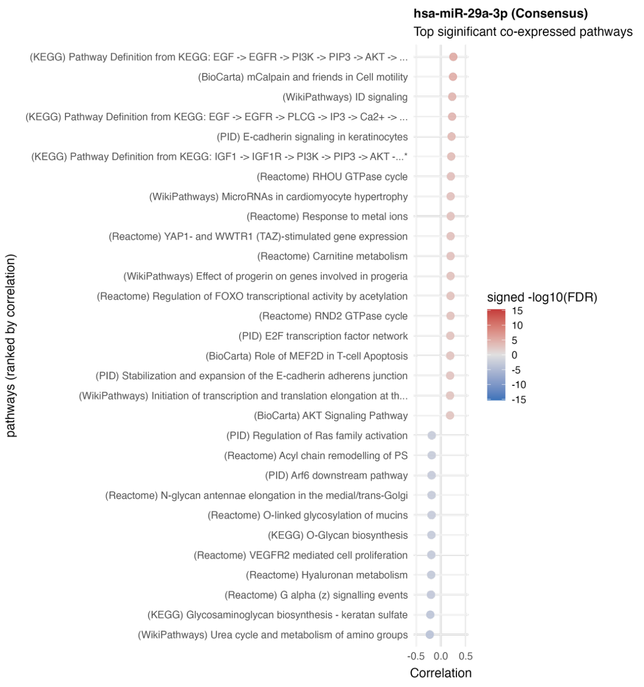

### Supplementary Figure 2.

**Cluster 1**

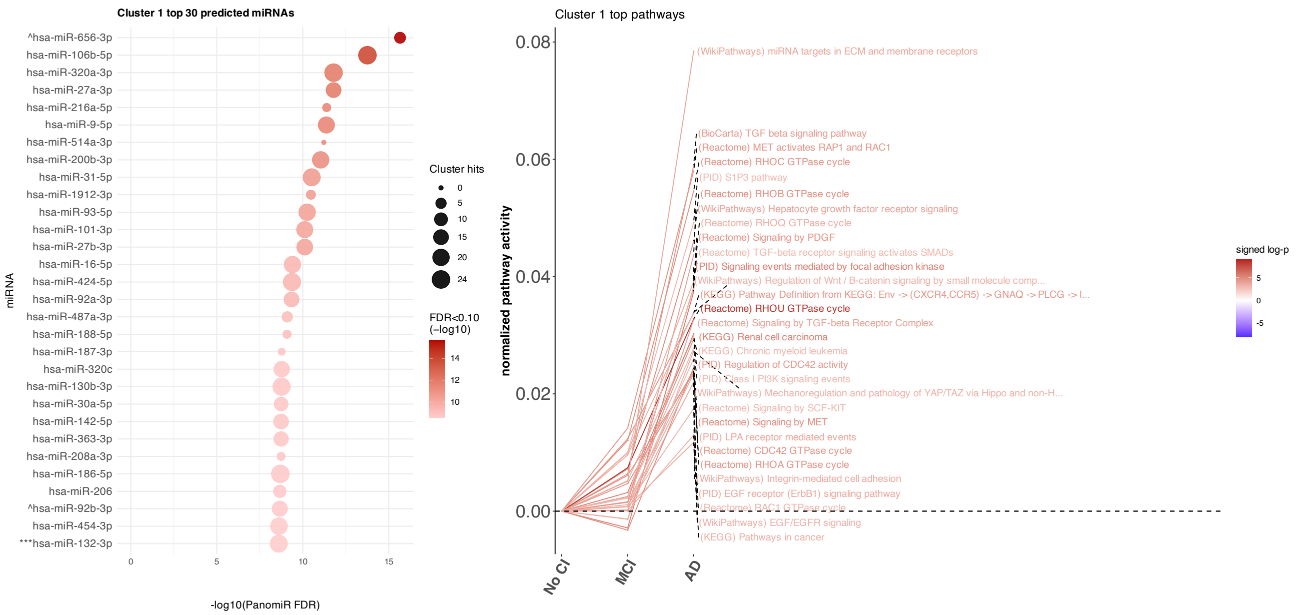


**Supplementary Figure 2. Characterization of cognitive-associated pathways across PanomiR clusters. Related to Figure 5.** The left panel shows the top 30 cognitive-associated pathways in each cluster. Y-axis represents the average activity of each pathway with levels of cognitive loss, centered on the baseline activity in subjects with no cognitive impairment. The right panel shows the top 30 miRNAs inferred by PanomiR that are enriched the cluster of pathways represented on the left panel. Node sizes are proportional to the number of individual pathways that are enriched in the targets of each miRNA. Tarbase V9 was used as the reference of miRNA-mRNA targeting events.

### Supplementary Figure 2 continued

**Cluster 2**

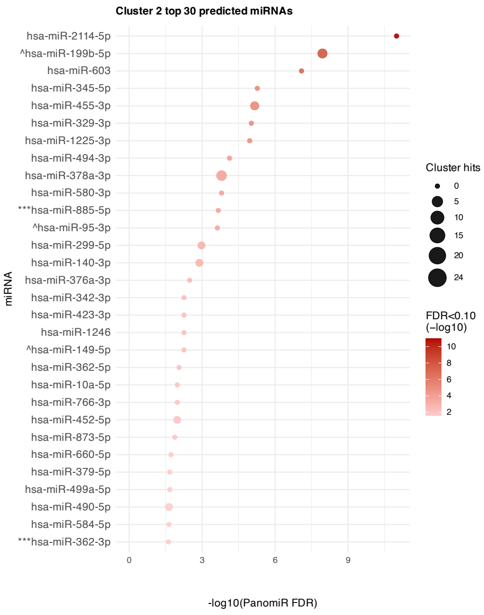


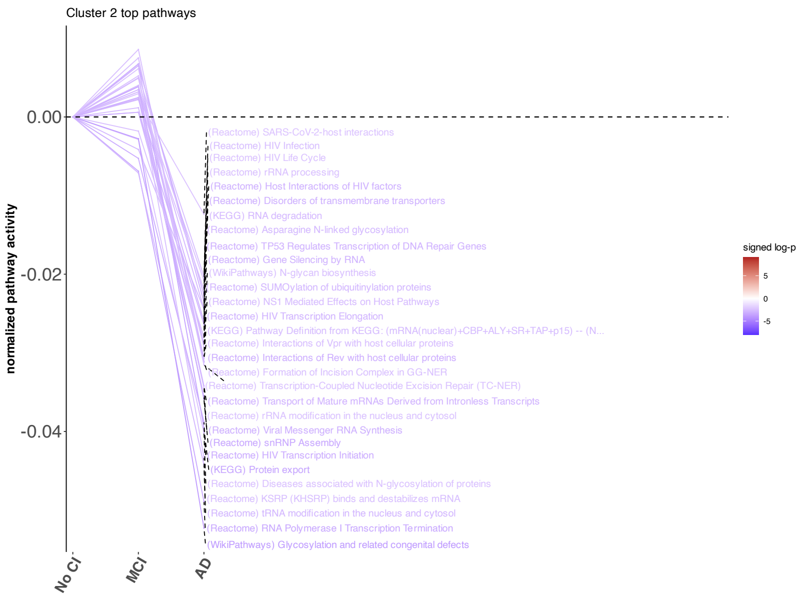

### Supplementary Figure 2 continued

**Cluster 3**

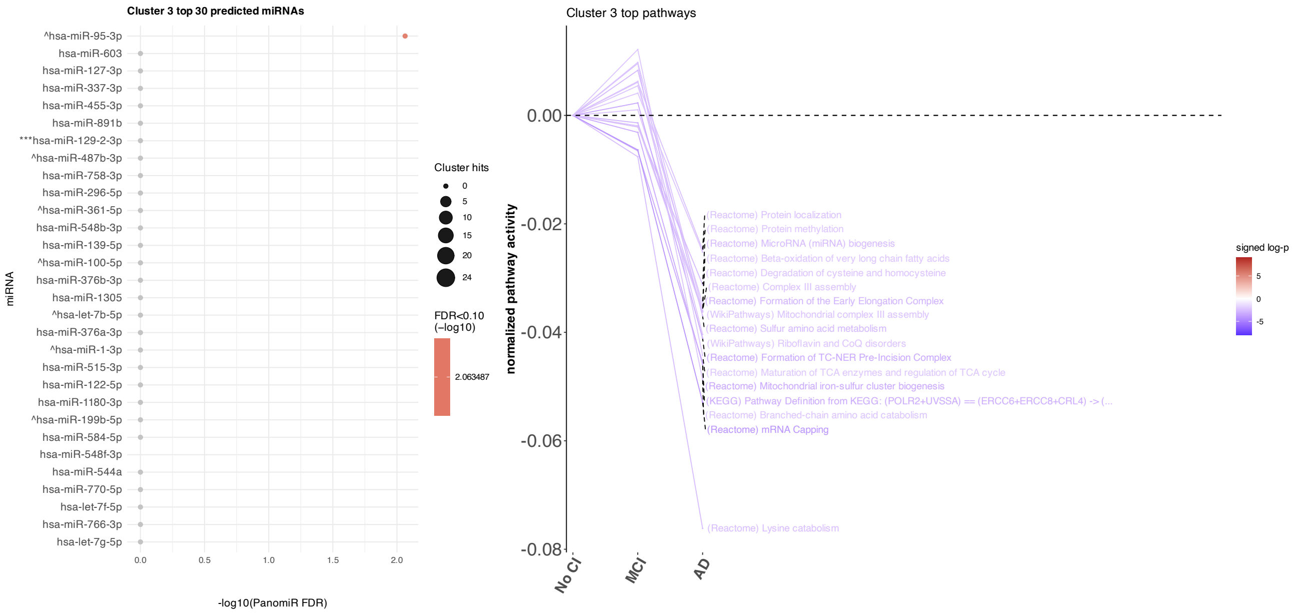

### Supplementary Figure 2 continued

**Cluster 4**

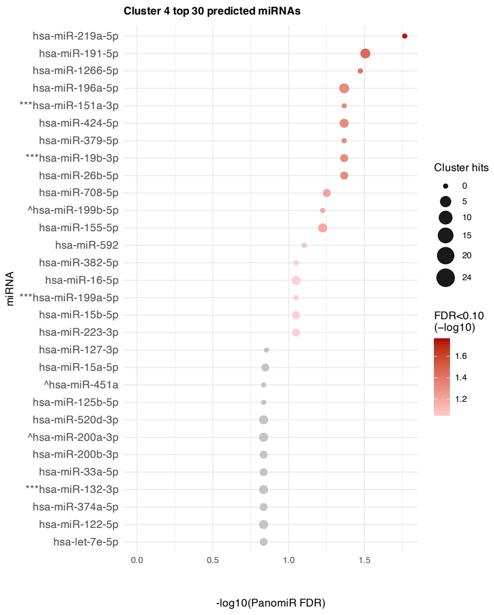


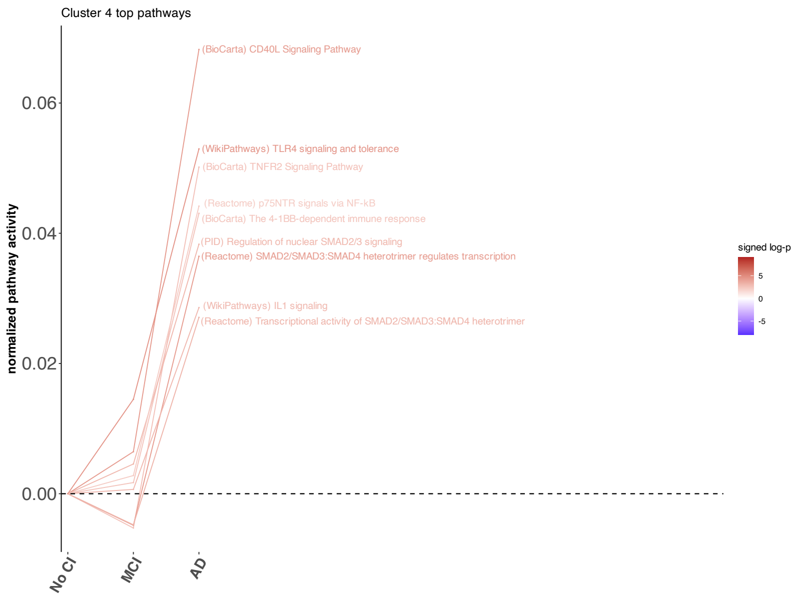

### Supplementary Figure 2 continued

**Cluster 5**

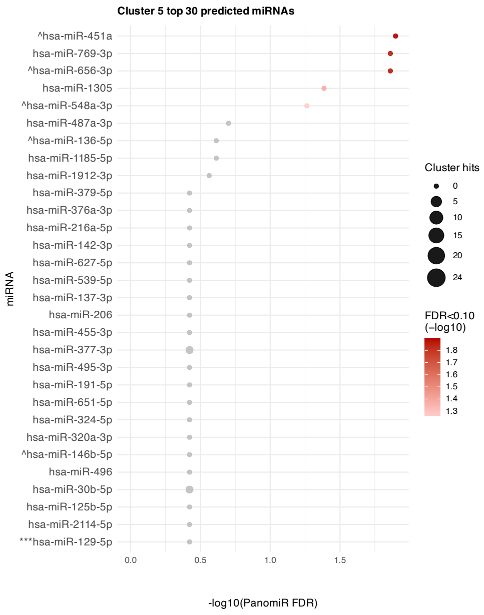


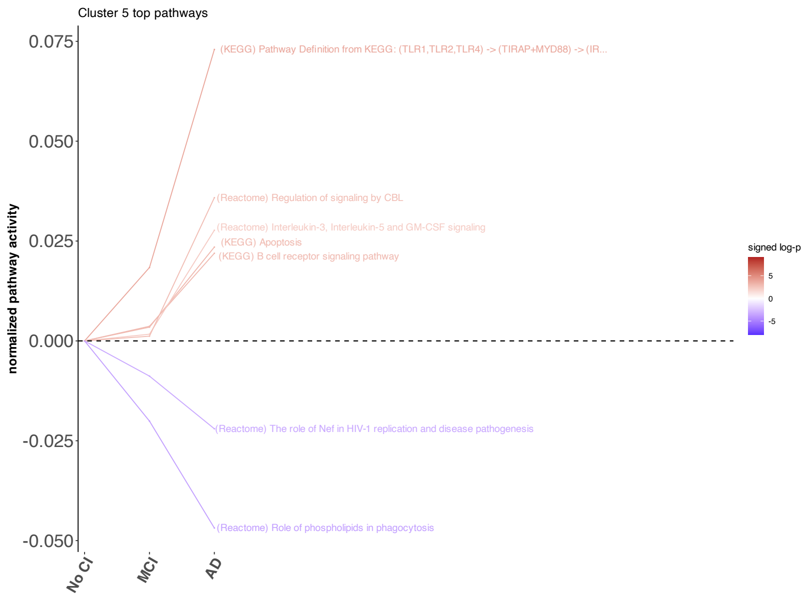

### Supplementary Figure 2 continued

**Cluster 6**

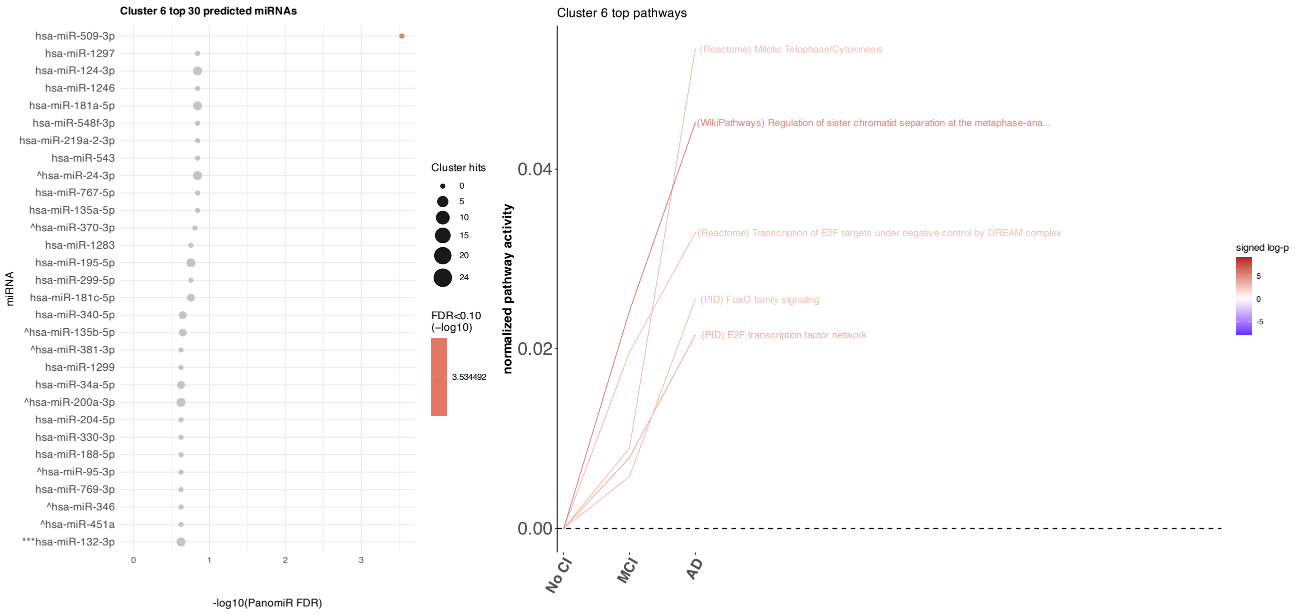

### Supplementary Figure 3.

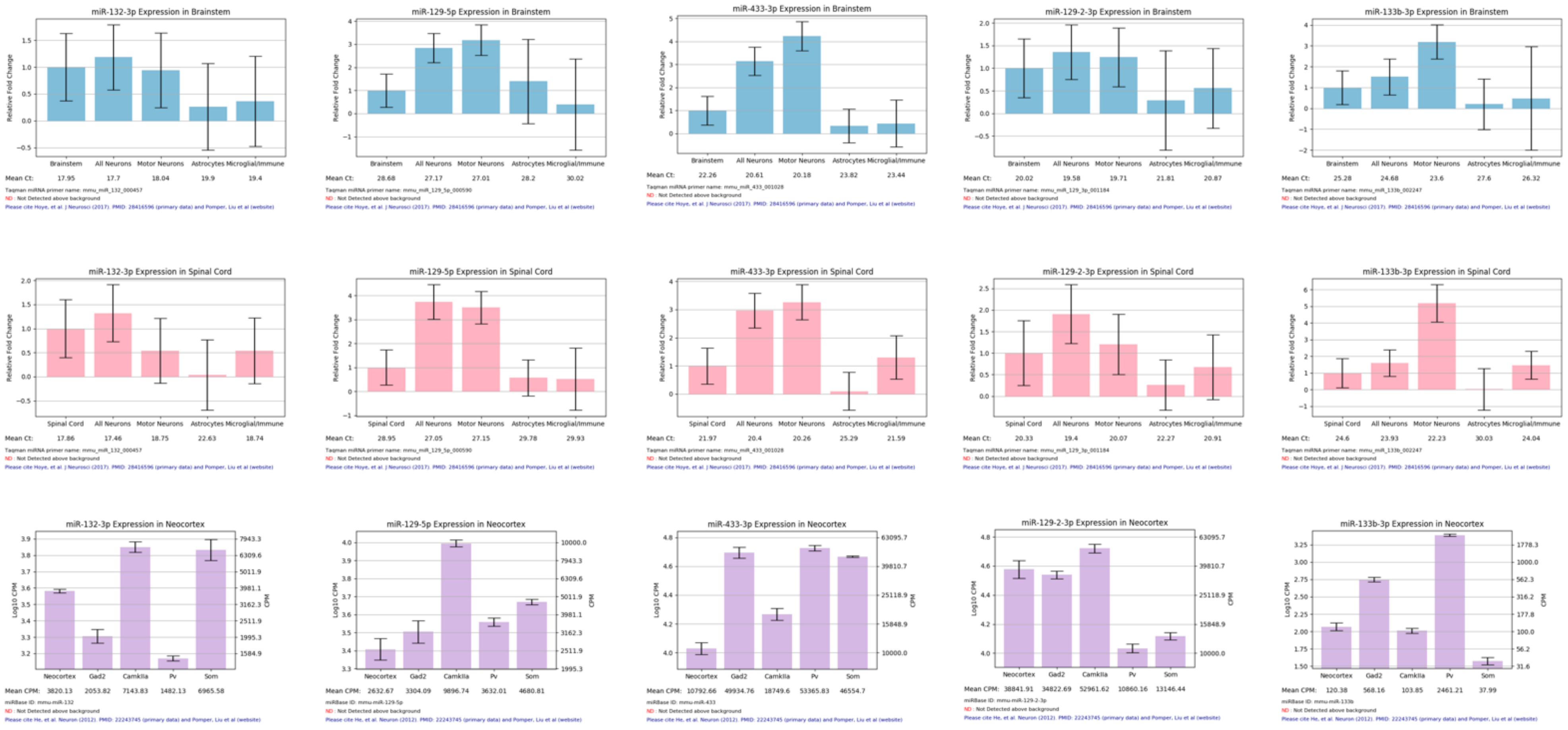


**Supplementary Figure 3. Cell-specific miRNA expression. Related to Figure 6 and Table 2. Cell-specific expression was retrieved from miRNA.wustl.edu. Panels represent miRNA expression in brain stem, spinal cord, and neocortex in mice.**

### Supplementary Figure 3 continued

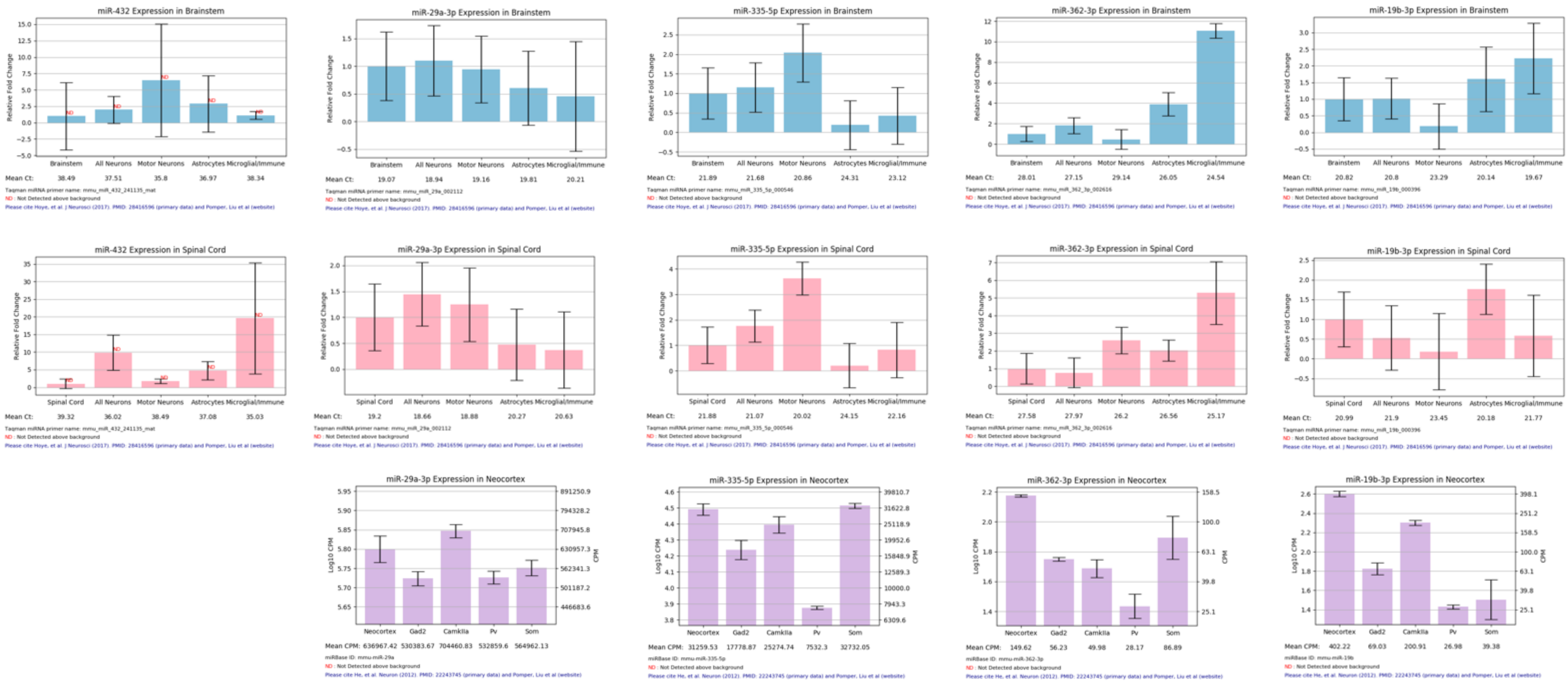

### Supplementary Figure 3 continued

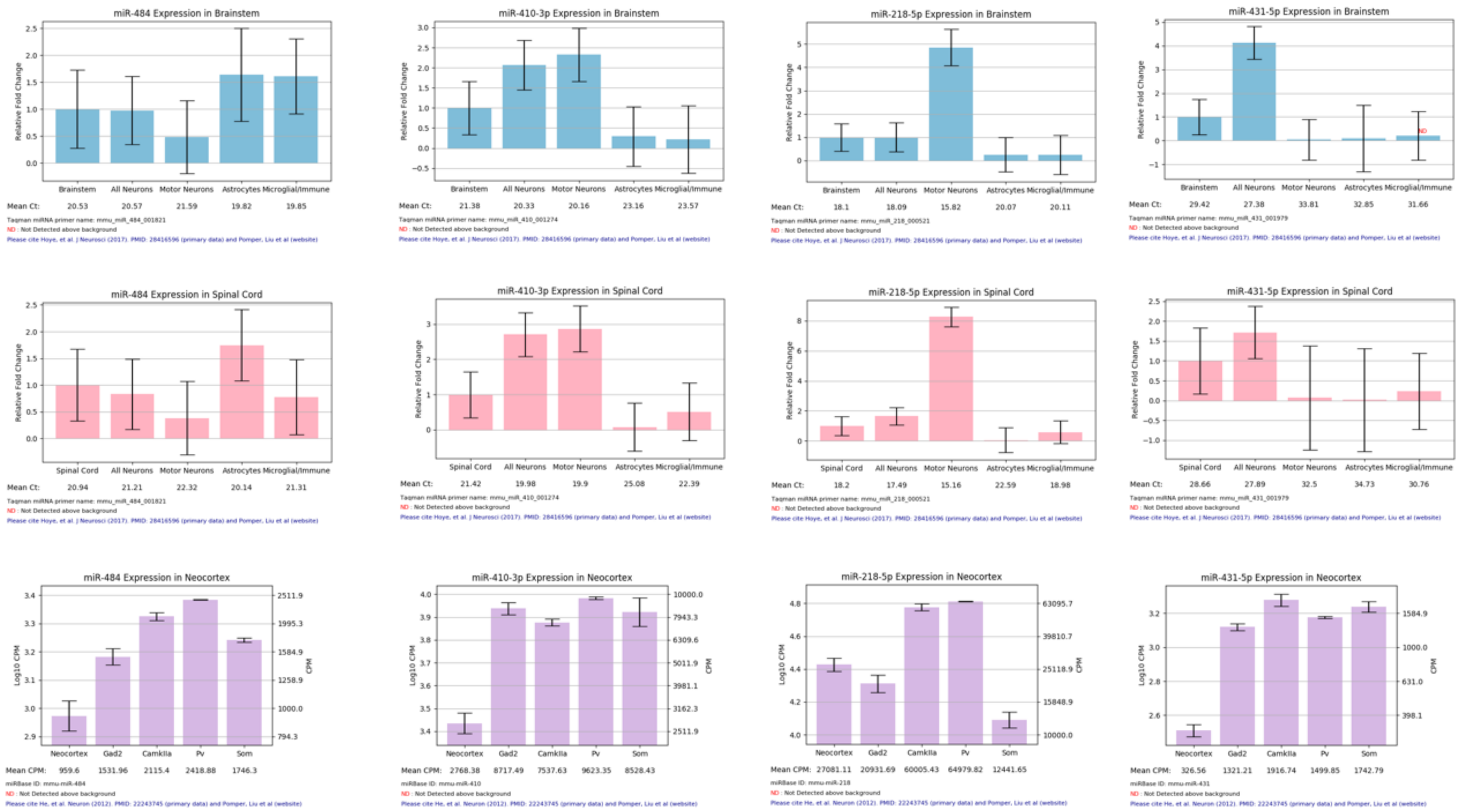
